# A single-cell atlas of ribosomal protein heterogeneity across human tissues reveals phenotypes of biological and clinical significance

**DOI:** 10.1101/2025.06.23.661225

**Authors:** Aishwarya Murali, Chartsiam Tipgomut, Anand D. Jeyasekharan, Himanshu Sinha

## Abstract

Ribosomes, once considered homogenous, exhibit dynamic compositional heterogeneity driven by differential ribosomal protein (RP) gene expression, modulating translational control. The exact cellular contributions to ribosomal heterogeneity in human tissues and their biological or clinical significance remain largely unknown. This study addresses these by mapping the expression of 76 cytoplasmic RP genes across 161 cell types from 15 human tissues using single-cell RNA sequencing from the Human Cell Atlas. We reveal extensive tissue- and cell-type-specific RP expression patterns, with *RPL23, RPS20, RPS17* and *RPL27A* showing variability across most tissues. The testis cells exhibited the greatest variability and the largest number of variable RP genes, with distinct RP signatures distinguishing germ and somatic lineages; these signatures form temporally coordinated expression modules throughout spermatogenesis. In the context of disease, a comparative analysis of male infertility patients revealed widespread RP gene dysregulation in testicular cell types, highlighting the importance of proper RP composition for reproductive health. Furthermore, given the link between RP gene mutations and inherited anaemia syndromes, we investigated RP gene expression during erythropoiesis. We observed disrupted RP gene expression in Diamond-Blackfan Anaemia patients, contrasting with stable patterns in normal erythropoiesis. Our findings underscore the underappreciated cellular specificity and dynamic regulation of RP gene expression, strongly implicating ribosome compositional heterogeneity as fundamental to both cellular identity and disease pathogenesis.

## INTRODUCTION

Ribosomes are key organelles made up of rRNA and ribosomal proteins (RPs) that translate mRNA into polypeptides essential for cellular functions, growth, and the development of the organism^1–3^. However, recent research has revealed that ribosomes are not uniform in terms of their protein composition but are dynamic and heterogeneous complexes that perform translation^4–7^. Ribosome heterogeneity and specialisation represent critical concepts in understanding the regulation of gene expression and protein synthesis. Heterogeneity arises through diverse mechanisms, including variations in rRNA^8,9^, differential stoichiometry of RP genes^4,10–12^ and their post-translational modification^13,14^. When such a heterogeneous ribosome drives translation of specific pools of transcripts, leading to a particular function or phenotype, it becomes a specialised ribosome^15–21^.

Functional specialisation of ribosomes has been observed in various organisms due to the absence or sub-stoichiometric levels of specific RP genes or proteins^10,12,17,22–25^. For example, changing one RP paralog (*RPL1B*) for another (*RPL1A*) reduced mitochondrial gene translation in yeast^26^. In fruit flies, the deletion of *RPS5B* led to defects in egg chamber formation in female germ cells^27^. Similarly, in mouse embryonic stem cells, heterogeneous ribosomes were found to cause selective translation of different pools of mRNA^11^.

Stoichiometric changes in RP gene expression are an essential aspect of ribosome heterogeneity and specialisation^23,28^. In humans, 79 different RPs combine with four rRNA molecules to form the large and small subunits of the ribosome. Each RP was found to be expressed differentially across various tissues, tumours and developmental stages in humans^23,29–31^. This differential expression can contribute to the formation of specialised ribosomes tailored to meet the specific translational needs of a particular tissue or a physiological condition^32–35^. Further, in humans, the functional effects of RP heterogeneity result in different pathologies^36–41^. RP genes were found to be mutated in ribosomopathies like Diamond-Blackfan Anaemia (DBA), resulting in defective ribosome biogenesis, leading to defective erythropoiesis^42^. Knockdown of *RPL35* in neuroblastoma tissues was found to affect HIF1α expression, affecting the proliferation of the NB cells^35^ and knockdown of specific RP genes was found to alter different gene subsets translationally in cancer cell lines^34^.

Using bulk transcriptome data, many studies have highlighted the tissue-specific gene expression of RP genes^23,29–31^. Bulk analyses average signals across diverse cell populations, which obscures important cell-type-specific RP gene expression patterns that could underlie specialised ribosomal functions within complex tissues. The single-cell RNA sequencing (scRNA-seq) data from the Human Cell Atlas (HCA) provides an extensive resource of single-cell transcriptomic profiles of various human tissues and cell types^43,44^, allowing one to investigate stoichiometric differences in RP expression across cell types and tissues (46–49). Does the tissue microenvironment influence changes in RP gene expression profiles, and do the various cell types within a tissue exhibit similar expression profiles? Do specific cell types maintain consistent RP gene expression profiles irrespective of the tissue microenvironment they inhabit? Further, do RP gene expression profiles change during cell type differentiation in a way similar to gene expression profile changes during embryogenesis^49–51?^

Using HCA data, we first extracted highly expressed RP gene expression profiles of all 161 constituent cell types of 15 tissues. Analysis of each RP gene expression variance across all the cell types showed that for a given RP gene, some tissues had high expression variability, while others exhibited little to no variation across the constituent cell types. This indicated that among the 15 tissues analysed, some tissues had all RP genes expressed uniformly across all cell types, while others displayed distinct expression profiles for each RP gene in different cell types. Using this framework, we revealed that in the testis, RP genes had expression heterogeneity in various stages of spermatogenesis, with some RP genes expressed higher than others in a stage-specific manner. Further, we illustrated that RP gene expression patterns vary among adult fertile, infertile and infant testes. Similarly, we uncovered dysregulation of RP gene expression patterns during erythropoiesis in DBA. This RP gene expression variability suggested that the regulation of RP genes was tightly controlled and context-dependent, which could have implications for cell type function and developmental stages.

## METHODS

### Data collection and processing

10x Genomics-based single-cell RNA sequencing data for 15 normal tissues and tissues from male infertility were obtained from the Human Cell Atlas (HCA) public repository (https://data.humancellatlas.org/). This data (Table S1) contained scRNAseq data from 14 tissues of a single donor (GSE159929)^52^, adult testis (GSE112013), infant testis (GSE120506)^53^, idiopathic non-obstructive azoospermia (NOA), retrograde oligozoospermia and Klinefelter syndrome (GSE169062)^54^. In addition, we analysed the normal and DBA bone marrow dataset (GSE222368)^55^ to study the RP gene expression changes in erythropoiesis.

R version 4.3.1 or later and the Seurat R package (v5.0.1)^56^ (https://satijalab.org/seurat/) were used for data analysis. Using the Merge function, we combined data from studies with multiple patients into a single Seurat object. To remove doublets and apoptotic cells, a filtering process was applied to the count matrix from each dataset based on the criteria established in the original studies, including thresholds for minimum and maximum genes per cell and the percentage of mitochondrial reads.

We employed ScTransform (v0.4.1)^57,58^ for normalisation because it effectively addressed technical variations arising from sequencing depth, particularly for highly expressed genes like RPs, compared to traditional log-normalisation methods. Dimensionality reduction was done using uniform manifold approximation and projection (UMAP)^59^, with the clusters formed using the Louvain algorithm using 50 principal components^60,61^. The elbow plot analysis indicated that the percentage of variance explained reached an early plateau across all tissues, with the most significant decline occurring within the first 10 to 20 principal components (PCs). However, to effectively capture biologically meaningful heterogeneity while reducing the technical noise inherent in single-cell data, we chose to use 50 PCs for downstream analysis. Harmony-based^62^ or CCA-based^63^ integration was applied for datasets involving multiple patients prior to clustering. The parameters used for preprocessing the Seurat objects are given in Table S1. Cell types were assigned based on marker gene information provided in the original publications and validated using the EnrichR web server^64^ and the Tabula Sapiens reference atlas^65^. The entire workflow of the analysis is described in Figure 1A.

**Figure 1.**
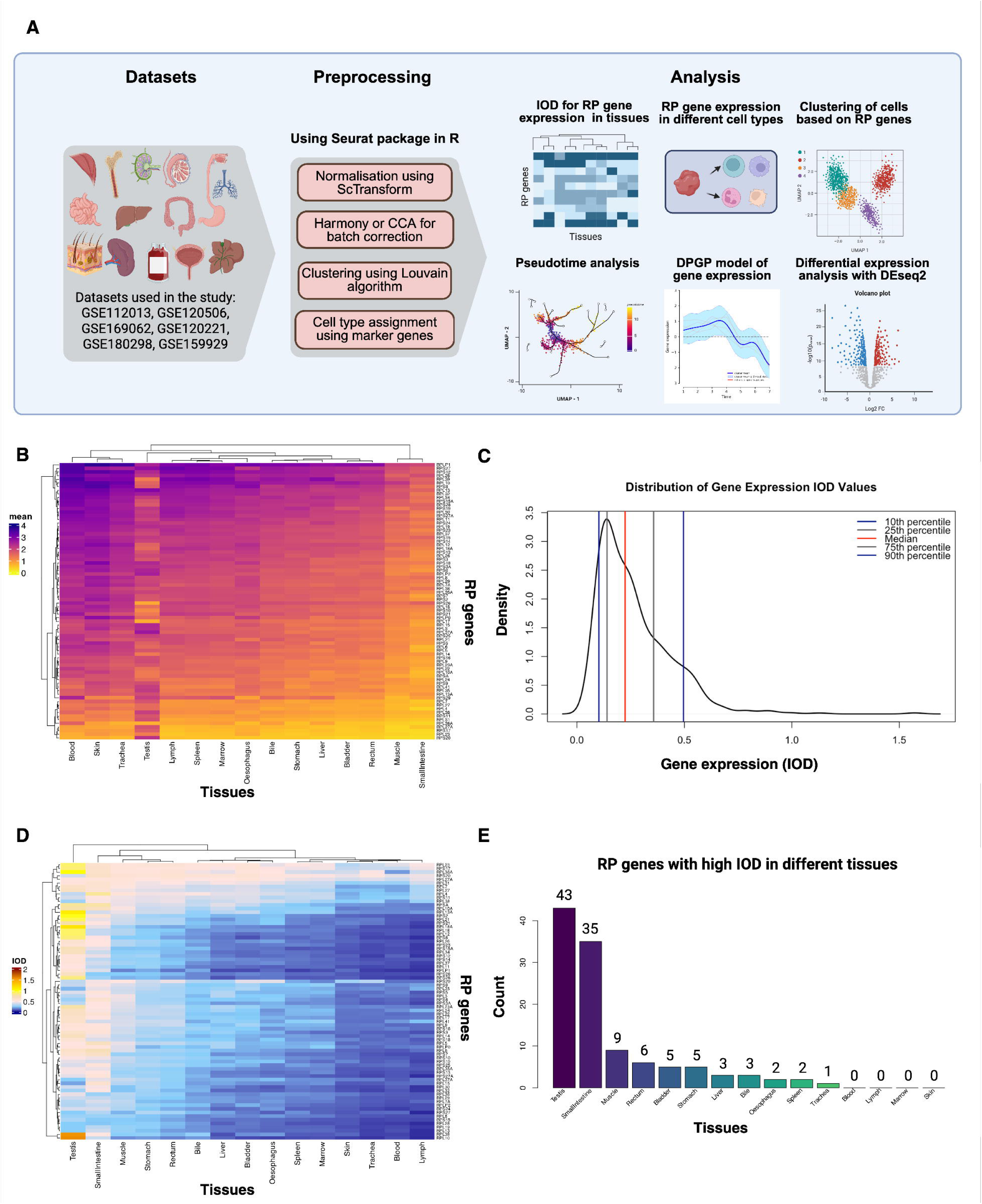
**Tissue-level expression patterns and variability of RP genes**. (A) Schematic overview of the scRNA-seq data analysis of RP genes from the Human Cell Atlas datasets. (B) Heatmap depicting the mean normalised expression of 76 cytoplasmic RP genes (rows) across 15 human tissue samples (columns). Tissue samples include blood, skin, trachea, marrow, spleen, oesophagus, bladder, liver, bile, stomach, rectum, muscle, small intestine, testis, and lymph. Colour intensity scale bar (0 to 4) corresponds to the average SCTransform-normalised expression level of RP genes within each tissue. Both genes and tissues are hierarchically clustered. (C) Density distribution of the IOD values of RP genes across the constituent cell types of the 15 tissues studied. The x-axis represents the IOD values. The y-axis represents the density of genes at a given IOD value. The coloured vertical lines within the density plot denote specific percentiles of the ID distribution: the 10th (blue), 25^th^(grey), 50^th^ (red), 75^th^ (grey), and 90^th^ (blue) percentiles. (D) Heatmap depicting IOD for RP gene expression within the 15 human tissue samples (scale 0 to 2). The y-axis represents the RP genes. The x-axis lists the tissue samples, clustered according to IOD values. (E) Histogram showing the number of RP genes that exhibit a high IOD value (above the 90^th^ percentile) in each tissue. The y-axis represents the count of RP genes having high IOD, and the x-axis lists different tissues. Created in https://BioRender.com.

### Comparison of RP gene variation across tissues and cell types

A total of 76 cytoplasmic RP genes were considered for analysis, excluding X and Y chromosome-linked (*RPS4X*, *RPS4Y1*, *RPS4Y2*). To assess tissue-level RP gene expression, we averaged the normalised expression of each RP gene across all cells within a specific tissue. We did not aggregate the data as the number of cells in each tissue differed. This allowed for a direct comparison of RP gene expression patterns between tissues. Since different tissues were found to have differences in their RP gene expression levels, to study the heterogeneity of RP genes within the tissues, we calculated the index of dispersion (IOD, variance/mean) to enable proper comparison. For this, in testis (with multiple donors), a single donor with the highest number of cells was used, as the other tissues were also from a single donor. The mean (Figure S1A) and IOD (Figure S1B) values were also checked across multiple donors for similarity of results (see Supplementary results). The IOD values were plotted as a distribution, and the values above the 90^th^ percentile were taken as high IOD, while the values below that were considered as low IOD (Figure 1C). A hierarchically clustered heatmap was generated to visualise the mean and variance of RP gene expression across all tissues. We analysed RP gene expression patterns across different cell types within these tissues to investigate the underlying causes of RP gene expression variability observed in specific tissues.

### Clustering of cells using RP genes as variables

A matrix was formed from the count data from the Seurat objects of each tissue for only RP genes and the highly variable genes without RP genes (referred to as the marker gene-based cluster). A new Seurat object was created for both the count matrices, and normalisation was done using ScTransform. After batch correction using Harmony, dimensionality reduction was performed using UMAP using 37 and 50 principal components, respectively. The clusters were formed using the Louvain algorithm with a resolution of 1. The visualisation in UMAP was done based on the cell types assigned to each cell in the previous section, and the RP-based clusters and marker gene-based clusters were compared.

### Identification of differentially expressed RP genes in constituent cell types of tissues

Using the Wilcoxon rank-sum test, the FindAllMarkers function in Seurat was employed to identify differentially expressed genes in each cell type. Genes exhibiting an average log fold change greater than 0.25 and detected in at least 10% of cells within a cell type were returned by the analysis as output. Using an average log_2_foldchange cutoff of 1 and an adjusted p-value (Bonferroni correction) less than 0.05, the significantly differentially expressed genes were identified. In the case of tissues with multiple donors, the FindConservedMarkers function was used to identify the common differentially expressed genes across donors. The minimum p-value returned by this function was corrected using Bonferroni correction. Further, the genes with fold change value greater than +1 or less than -1 consistently in all donors and the adjusted p-value less than 0.05 were chosen. Volcano plots were drawn to find the significantly differentially expressed genes comparing one cell type with others.

### Pseudotime series analysis

The bone marrow dataset (GSE222368) was analysed only for three normal and two DBA cases to identify the erythrocytes and progenitor cells based on *HBA1, CA1, SLC4A1, GYPA, TFRC, HMGA1* and *KIT* expression. This dataset encompassed both normal individuals (n = 3) and individuals with Diamond-Blackfan Anaemia (DBA, n = 2). These erythrocytes and progenitors for all the individuals were converted to a cell dataset object using Monocle3^66,67^. Progenitor cells with high KIT expression were defined as the pseudotime origin^55^. After this, the data was subset to different patients and healthy donors, and the pseudotime trajectory was divided into equal bins and used for the Dirichlet Process Gaussian Process (DPGP) analysis. The bins were defined based on the pseudotime values of normal erythropoiesis and defined as stages 1 to 7. A similar pipeline was followed for all the normal donors and patients of DBA. We also compared *GATA1* and *SPI1* expression with *RPS19* in all three cases^68,69^. Similarly, for spermatogenesis, the spermatogonial stem cells were used as a starting point, and the cluster order and pseudotime values were compared to ascertain the correctness of the order of cell types being assigned.

### Dirichlet process Gaussian process model to identify RP-based clusters in differentiation processes

To cluster the RP genes with similar expression patterns in spermatogenesis and erythropoiesis, we used the DPGP model, a nonparametric Bayesian method where a Gaussian process models the temporal gene expression trajectories while the Dirichlet process identifies the optimal number of clusters^70^. The normalised RP gene expression in different cell types of spermatogenesis and stages of erythropoiesis (assigned based on pseudotime) was averaged and used as input for DPGP analysis.

### Identification of differentially expressed genes between normal and pathological testes

RP genes that are differentially expressed between normal and pathological testis were identified using DESeq analysis. Gene expression was aggregated for each cell type for each donor or patient to create a pseudobulk expression matrix, and the differentially expressed genes were identified using DESeq2^71^. Batch effects and variations in aggregated expression due to different cell numbers per sample were accounted for in the DESeq2 model.

### Analysis of common cell types across tissues

From every tissue, the common cell types were identified. The Seurat objects were subset for the cell types, and these subsets were merged. For this analysis, only those cell types from the GSE159929 dataset were used as they belonged to the same individual. The merged Seurat objects were subjected to ScTransform normalisation again, and the medians of all tissues were centred for differential expression analysis using PrepSCTFindMarkers. Further, PCA was done, and using 50 PCs, the tissues were clustered using the Louvian algorithm. Further, using the Wilcox rank sum test (FindMarkers), the differentially expressed genes between the tissues were identified.

## RESULTS

### RP genes exhibit tissue-specific mean expression levels

In this study, we extracted and considered the normalised RP gene expression of 76 cytoplasmic somatic RP genes in humans. In each of the scRNA-seq datasets of 15 tissues from non-diseased individuals from the HCA database, we averaged gene expression of all the constituent cells for a given RP gene (Table S1). The average RP gene expression value accounted for varying cell numbers across tissues and allowed direct comparisons.

The hierarchical clustering of the average RP gene expression of the constituent cells of a tissue showed that each RP gene had varying expression patterns across tissues (Figure 1B). For example, *RPLP1, RPS27, RPS12* and *RPL28* had high expression in most tissues but a comparatively lower expression in the small intestine and muscle. In contrast*, RPLP0* and *RPL17* had variable expression across tissues, while *RPL36A* consistently showed low expression across all tissues except blood (Figure 1B). It was also observed that RP gene expression was heterogeneous within a tissue, i.e., all RP genes were not uniformly expressed as expected. The mean expression values of RP genes across tissues are summarised in Table S2. These results highlight the tissue-specific patterns of RP gene expression, which were similar to those observed in the previous studies ^31,72^. This analysis revealed considerable heterogeneity in RP gene expression within and across tissues (Figure 1B). We analysed RP gene expression variation across constituent cell types within a tissue to further explore this expression heterogeneity.

### Cell-to-cell variability defines RP gene expression within tissues

In the previous section, we observed that average gene expression values for RP genes within a tissue varied. Given that each tissue consisted of different cell types and the scRNA-seq dataset had gene expression values of multiple cells of each cell type, we wanted to explore how RP gene expression values varied within a tissue. To analyse this, we calculated the index of dispersion (IOD) for each RP gene for all the cells within a tissue. A higher IOD value (IOD above 90^th^ percentile, Figure 1C) would indicate that the RP gene expression value varied across different cells within a tissue. An RP gene with a higher IOD value would have more expression variation in a tissue compared to an RP gene with a lower IOD value.

Hierarchical clustering of the IOD values of RP genes across tissues showed that tissues like the testis and small intestine had the highest RP gene expression variation and a greater number of RP genes with high IOD values across their cells (Figure 1D, 1E). Some tissues, like skin, trachea, blood, marrow and lymph, had the lowest IOD values, indicating that RP gene expression does not vary across the cells in these tissues. The patterns of IOD values revealed that RP genes such as *RPL39 and RPL10* had the highest IOD values in the testis, but were uniformly expressed (low IOD value) in all other tissues (Figure 1D). It was also observed that some RP genes, such as *RPL23, RPS20, RPS17* and *RPL27A*, had uniformly higher IOD values across most of the tissues, indicating that these genes were variable in most of the tissues tested. Further, gene *RPL4* was only variable in the small intestine and muscle, while it had a low IOD in other tissues. Finally, about 20 RP genes, including *RPL19, RPS15, RPL28,* and *RPS27*, had low IOD values in all tissues studied, indicating uniform expression in cells across tissues. These IOD results showed that the RP gene expression patterns vary considerably across the cell types of certain tissues. The IOD values of RP gene expression across tissues are summarised in Table S3. Further, we compared the IOD values of RP genes with those of random gene sets and ubiquitous gene sets. We found in testis and a few other tissues that RP genes had significantly different IOD values than other gene sets (see Supplementary results).

The variation in RP gene expression among cell types in different tissues highlighted the need to study specific cells for a better understanding of RP heterogeneity. To understand what drives high IOD values of RP genes across constituent cell types within a tissue, we analysed the expression of the RP genes within each cell type in a tissue. Cell identity was assigned using cell-specific marker genes (Table S4). For the 14 tissues and their constituent 150 cell types, RP genes were found to be differentially expressed in certain cell types (Table S5). The comprehensive list of RP gene expression upregulated and downregulated in the cell types of different tissues is given in Table 1. Of the tissues with top IOD values, we found that both the testis and the small intestine had differences in RP gene expression among their constituent cell types. However, we did not proceed with the small intestine, as in addition to cell-type-specific changes, the lower mean of RP genes could have contributed to higher IOD. Further, we analysed RP gene expression for common cell types, namely fibroblasts, monocytes, CD8T, plasma, smooth muscle, NKT, B and T cells across the 14 tissues, excluding testis. We found that only a few of the cell types, like plasma cells of the stomach (Figure 2A), spleen (Figure 2B), endothelial (Figure 2C) and smooth muscle cells (Figure 2D) of the muscle, had tissue-specific RP gene expression. Further, a few of the cell types had a small number of RPs that were differentially expressed in a tissue-specific manner (Table 2, S5).

**Figure 2.**
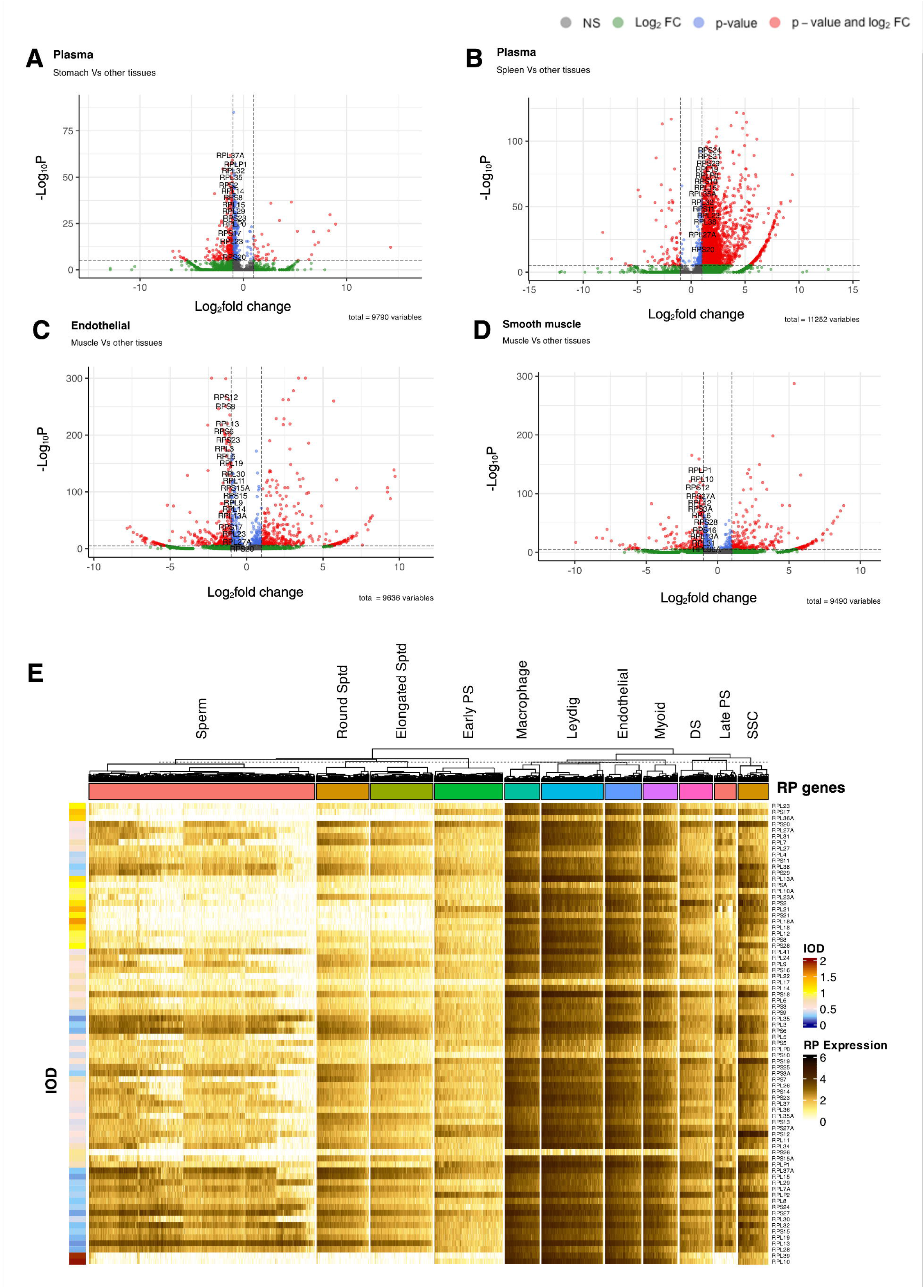
**Cell-type-specific expression and variability of RP genes**. (A) A volcano plot illustrating the differential expression of genes in plasma cells from stomach tissue compared with the same cell type in other tissues. The x-axis represents log_2_ fold change, and the y-axis represents -log_10_ (p-value). Genes are coloured based on statistical significance: NS (not significant, grey), significant by log_2_ fold change only (green), significant by p-value only (blue), and significant by both p-value and log_2_ fold change (red). Specific RP genes (e.g., *RPL37A*, *RPLP1*) that are significant by both p-value and log_2_ fold change are labelled. (B) A volcano plot illustrates the differential expression of genes in plasma cells from spleen tissue compared with the same cell type in other tissues. Axes, colouring, and labelling conventions are similar to panel (A). A specific RP gene (e.g., *RPS24, RPS21*) that is significant by both p-value and log_2_ fold change is labelled. (C) A volcano plot illustrates the differential expression of genes in endothelial cells from muscle tissue compared with the same cell type in other tissues. Axes, colouring, and labelling conventions are similar to panel (A). A specific RP gene (e.g., *RPS12, RPS8*) that is significant by both p-value and log_2_ fold change is labelled. (D) Volcano plot illustrates the differential expression of genes in smooth muscle cells from muscle tissue compared with the same cell type in other tissues. Axes, colouring, and labelling conventions are similar to panel (A). A specific RP gene (e.g., *RPLP1, RPL10*) that is significant by both p-value and log_2_ fold change is labelled. (E) Heatmap displaying the normalised expression of 76 RP genes (rows) across major testicular cell types (columns): Sperm, Round Spermatids (Round Sptd), Elongated Spermatids (Elongated Sptd), Early Primary Spermatocytes (Early PS), Macrophages, Leydig cells, Endothelial cells, Myoid cells, Differentiating Spermatogonia (DS), Late Primary Spermatocytes (Late PS), and Spermatogonial Stem Cells (SSC). Colour intensity scale (0 to 6) within the heatmap indicates the SCTransform-normalised expression level of each RP gene in each cell type. A colour bar to the left of the heatmap (scale bar: 0 to 2) indicates IOD for each RP gene across these testicular cell types. Columns (cell types) are hierarchically clustered. Cell types are grouped by coloured bars at the top. Created in https://BioRender.com.

**Table 1.**
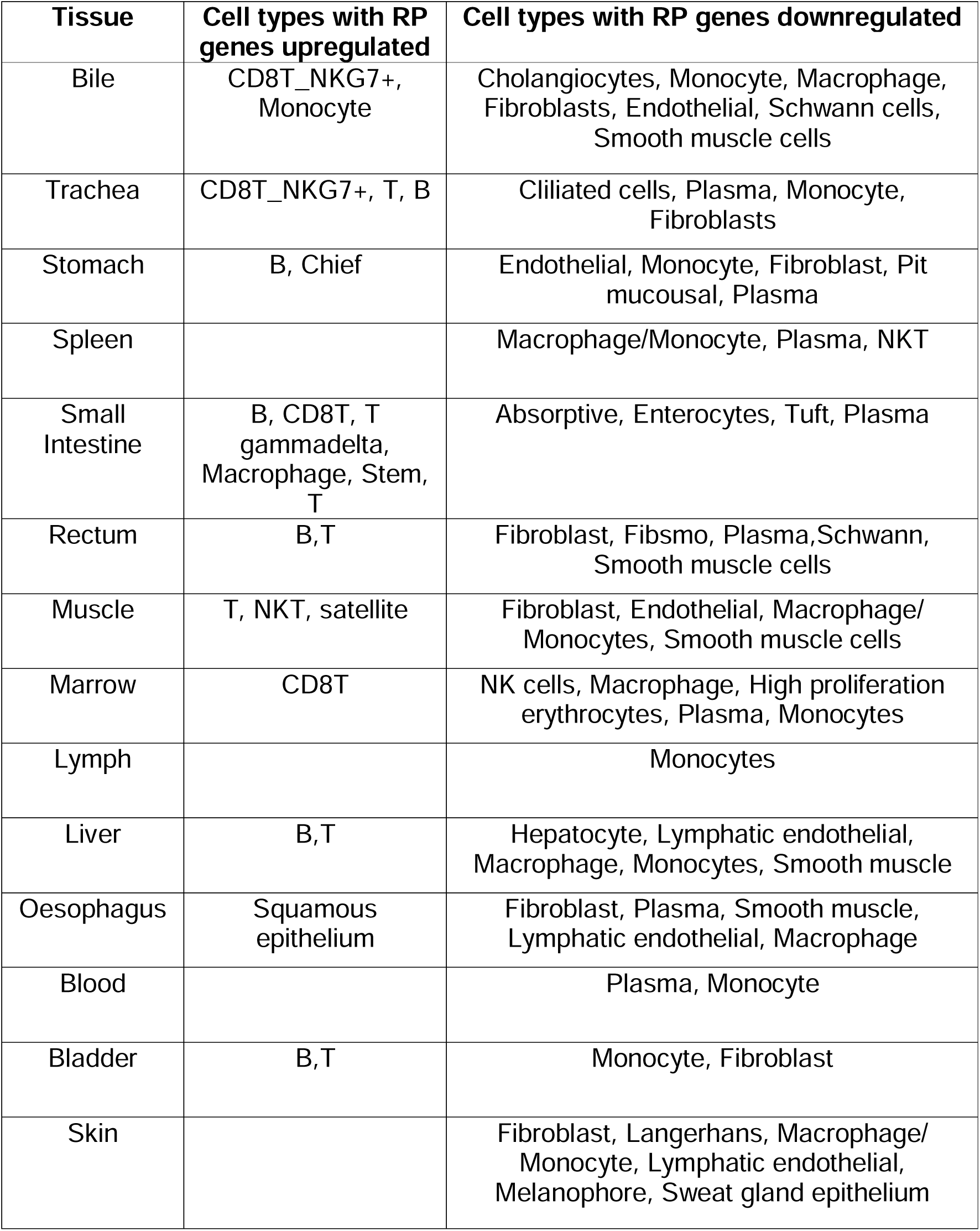
Summary of cell types with differential ribosomal protein gene expression in various tissues.

| Tissue | Cell types with RP genes upregulated | Cell types with RP genes downregulated |
| --- | --- | --- |
| Bile | CD8T_NKG7+, Monocyte | Cholangiocytes, Monocyte, Macrophage, Fibroblasts, Endothelial, Schwann cells, Smooth muscle cells |
| Trachea | CD8T_NKG7+, T, B | Ciliated cells, Plasma, Monocyte, Fibroblasts |
| Stomach | B, Chief | Endothelial, Monocyte, Fibroblast, Pit mucousal, Plasma |
| Spleen |  | Macrophage/Monocyte, Plasma, NKT |
| Small Intestine | B, CD8T, T gammadelta, Macrophage, Stem, T | Absorptive, Enterocytes, Tuft, Plasma |
| Rectum | B,T | Fibroblast, Fibsmo, Plasma, Schwann, Smooth muscle cells |
| Muscle | T, NKT, satellite | Fibroblast, Endothelial, Macrophage/ Monocytes, Smooth muscle cells |
| Marrow | CD8T | NK cells, Macrophage, High proliferation erythrocytes, Plasma, Monocytes |
| Lymph |  | Monocytes |
| Liver | B,T | Hepatocyte, Lymphatic endothelial, Macrophage, Monocytes, Smooth muscle |
| Oesophagus | Squamous epithelium | Fibroblast, Plasma, Smooth muscle, Lymphatic endothelial, Macrophage |
| Blood |  | Plasma, Monocyte |
| Bladder | B,T | Monocyte, Fibroblast |
| Skin |  | Fibroblast, Langerhans, Macrophage/ Monocyte, Lymphatic endothelial, Melanophore, Sweat gland epithelium |

**Table 2.**
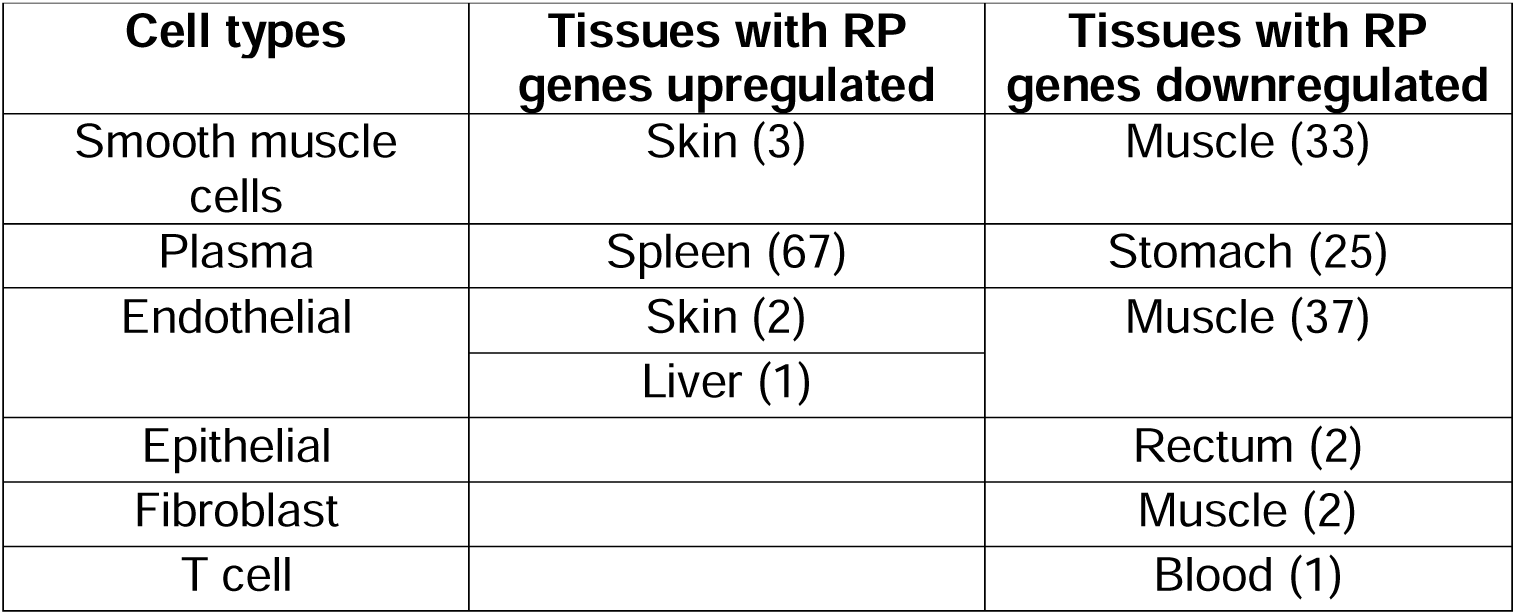
Summary of tissues with specific RP gene expression for common cell types (number of RP genes mentioned in the bracket)

### Distinct RP gene signatures define testicular germ and somatic cell lineages

Given that testis exhibited the highest IOD in our previous analysis, we next examined RP gene expression levels across all constituent somatic and germ cell types. It was observed that somatic cells, namely, macrophages, myoid cells, Leydig cells, and endothelial cells, had uniform gene expression for all the RP genes, with some RP genes having higher gene expression and others lower (Figure 2E). The germ cells, namely, spermatogonial stem cells, differentiating spermatogonia, early and late primary spermatocytes, round and elongated spermatids, and sperm, had differential RP gene expression (Figure 2E). This was true for the three independent donor samples (Supplementary results, Figure S1,S2, Table S6). Comparison of somatic and germ cell clusters of the testis using two features, highly variable genes (without RP genes) and RP genes, showed that germ cells formed similar clusters for both features, whereas somatic cells clustered differently for the two features (Figure 3A,3B). This indicated that RP gene expression was cell-type specific only for germ cells and not for somatic cells of the testis.

**Figure 3.**
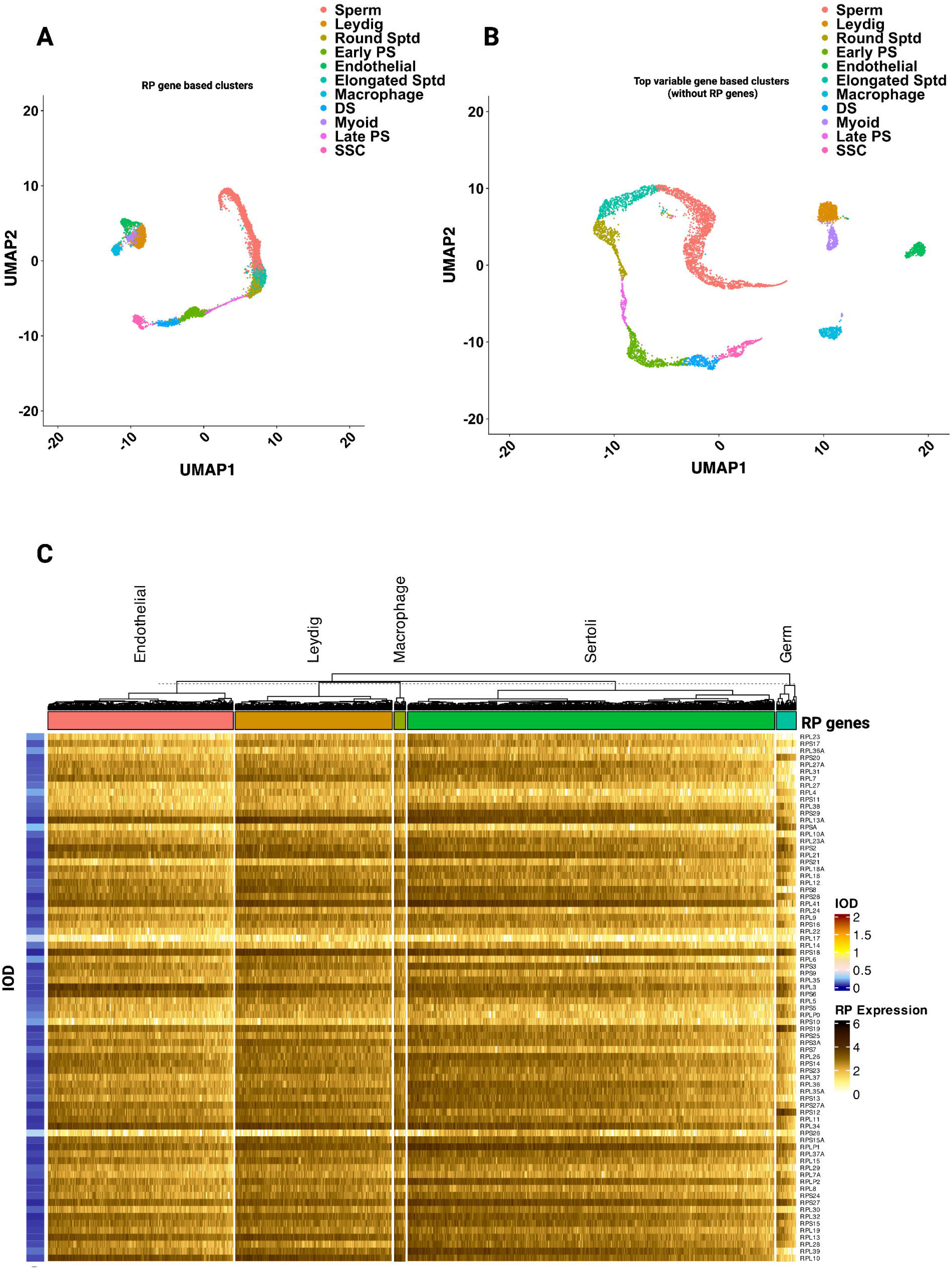
**Distinct RP gene expression signatures characterise specific cell types within the human testis**. (A) UMAP visualisation of single cells from human testis, clustered based solely on the expression profiles of 76 cytoplasmic RP genes. Distinct cell populations are resolved, including various stages of germ cell development (Sperm, Round Spermatids (Round Sptd), Elongated Spermatids (Elongated Sptd), Early Primary Spermatocytes (Early PS), Late Primary Spermatocytes (Late PS), Spermatogonial Stem Cells (SSC)) and somatic cell types (Leydig cells, Endothelial cells, Macrophages, Myoid cells). Each point represents a single cell, coloured by its assigned cell type. (B) Comparative UMAP visualisation of the same testis single cells, with clustering based on the expression of all other top 2,000 highly variable genes, excluding RP genes. Similar to panel (A), the legend indicates the specific cell types corresponding to the colours. (C) Heatmap displaying the normalised expression levels of individual RP genes (rows) across the different cell types identified within the infant testis (columns: Germ cells, Sertoli cells, Endothelial cells, Leydig cells, and Macrophages). The colour intensity represents the relative expression level of each RP gene (red for higher expression, blue for lower expression). Alongside the heatmap, a bar plot shows the IOD for each RP gene, indicating the variability of its expression across these testicular cell types. Cell types are grouped by coloured bars at the top. Created in https://BioRender.com.

To understand if the somatic cells and naïve and quiescent states of germ cells in infants were differentially expressing RP genes compared to the adult somatic and germ cells, we analysed scRNA-seq data from two 13-month-old infants (Figure 3C). As expected, infants’ testes had a higher representation of somatic cells, namely Sertoli, Leydig, endothelial cells, and macrophages, and a small number of naïve germ cells. Similar to what was observed for the adult testes’ somatic cells, the gene expression in infant testes across different cell types was also uniform for a given RP gene, with some RP genes having high expression and others having low. Compared to the infant somatic cells, for one type of naïve germ cells, the gene expression values were different for different RP genes. Interestingly, in the infant, only RPSA (p-value = 4.11E-08) was upregulated in germ cells, while 27 RP genes were downregulated compared to the somatic cells (average Wilcox rank sum test p-value = 0.00120, Table S7).

### Temporal clustering reveals dynamic RP gene expression modules during spermatogenesis

In adults, male germ cell development is a complex pathway through a series of states involving different cell proliferation and differentiation stages to generate sperm through spermatogenesis continuously. Given the differential expression of RP genes in various germ cells, we aimed to investigate whether the expression of RP genes followed a pattern that aligned with the distinct stages of spermatogenesis.

Using gene expression data from all germ cell genes, a pseudotime trajectory was identified. This pseudotime line was aligned with the cell-type-based order of cell differentiation during spermatogenesis. To analyse RP gene expression from discrete cells during the spermatogenesis process, we utilised a nonparametric Bayesian approach known as the DPGP model^70^. This method identified a longitudinal cluster of RP genes having similar patterns. This clustering allowed us to identify RP gene sets co-expressed during different stages of spermatogenesis (Figure 4 A-E).

**Figure 4.**
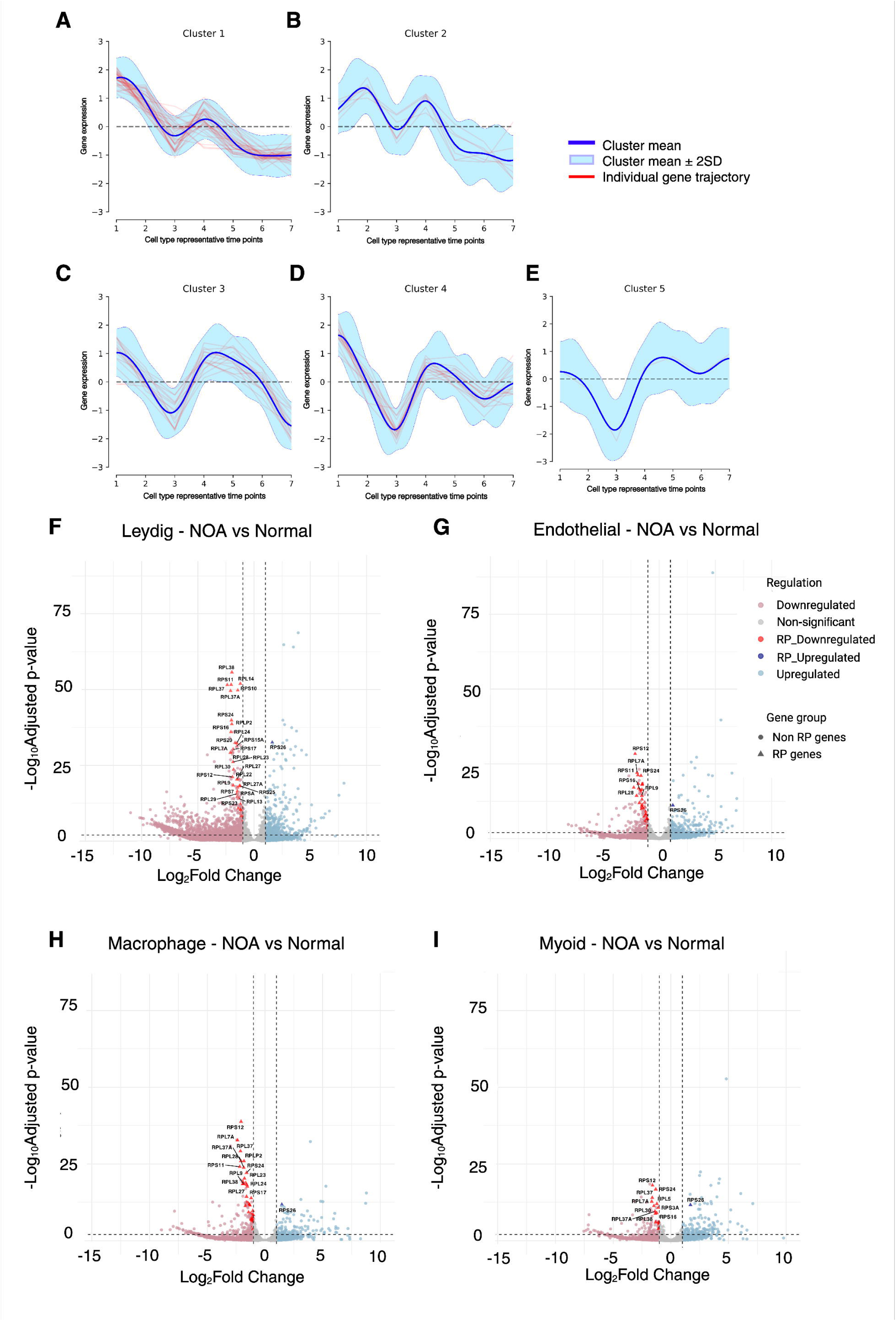
**Cell-type-specific dysregulation and clustered expression patterns of RP genes in male fertility**. (A-E) The DPGP model-based trajectories of RP gene expression profiles along spermatogenesis (x-axis) of five distinct gene clusters (Clusters 1-5). The y-axis represents the log-normalised ScTransform corrected gene expression values of RP genes. The x-axis represents the cell-type states of spermatogenesis at different representative time points (1-7): Spermatogonial stem cells (1), differentiating spermatogonia (2), early primary spermatocytes (3), late primary spermatocytes (4), round spermatids (5), elongated spermatids (6), sperms (7). Each small plot displays the normalised expression (y-axis) of individual RP genes within that cluster (red lines) as a function of pseudotime (x-axis). The “light blue zone” along each line in the plots represents the standard deviation (SD) of gene expression for the individual RP genes within that cluster, around the mean expression profile (blue line). (F-I) Volcano plots demonstrating differential gene expression in the pseudobulk aggregation in testicular somatic cell types when comparing samples from individuals with primary infertility to normal controls: (F) Leydig cells, (G) Endothelial cells, (H) Macrophages, (I) Myoid cells. In these plots, each point signifies a gene. The x-axis displays the log_2_ fold change, while the y-axis represents the -log_10_ (p-value). Genes showing statistically significant upregulation or downregulation are highlighted, with RP genes that are significantly upregulated “RP_Upregulated” (dark blue), RP genes that are significantly downregulated “RP_Downregulated” (dark red), Other, non-RP genes that are significantly upregulated “Upregulated” (light blue), Other, non-RP genes that are significantly downregulated “Downregulated” (light red), Non-significant genes “Non-significant” (grey). RP genes are indicated as triangles, and all other genes are shown as circles. Created in https://BioRender.com.

The DPGP analysis clustered RP gene expression into five clusters (Figure 4A-E). Cluster 1 (Figure 4A), which had 35 of 76 RP genes being considered, had an average gene expression decreasing from spermatogonial stem cells to the sperm stage. Whereas, Clusters 2 (Figure 4B) and 3 (Figure 4C) had RP gene expression patterns that changed with different stages of spermatogenesis. Interestingly, in Clusters 4 (Figure 4D) and 5 (Figure 4E), the RP gene expression patterns varied over the developmental timeline, for these RP genes and their expression increased from the spermatid stage to the sperm stage. In cluster 4, of the 17 RP genes, two RP genes, *RPL3* and *RPS6*, were observed to be upregulated in the sperm of humans^73^.

These patterns of RP gene clusters were largely maintained even when data from individual donors were analysed (Figure S3). The RP genes belonging to different clusters in the overall and specific donor samples are summarised in Table S8. These results indicated that the RP genes showed specific gene expression patterns during spermatogenesis, and they might play a role in individual stages of spermatogenesis.

### Cell-type-specific downregulation of RP genes in the testis associates with male infertility

The most severe form of male infertility, NOA, is a medical condition characterised by the complete absence of sperm in the ejaculate due to impaired or failed sperm production in the testes. We have shown above that RP gene expression varied across different germline stages but was uniform in somatic cells. Since there were no germ cells in NOA, we compared RP gene expression in the somatic cells of normal and NOA testes. Somatic cells are vital for enabling spermatogenesis, as they offer structural support, regulatory factors, and help facilitate the essential interactions between cells that are crucial for the development of germ cells^74,75^. Compared to the somatic cells from a normal testis, *RPS26* was found to be upregulated, and 19 RP genes were downregulated in somatic cells, namely, Leydig (Figure 4F) endothelial (Figure 4G), macrophages (Figure 4H) and myoid cells (Figure 4I) of the NOA testis (Figure 4F-I, Table S9).

Furthermore, gene ontology analysis of the differentially expressed genes between normal and NOA testes (Table S10) revealed that, in all the cell types of NOA, most RP genes were uniformly downregulated, while genes of different pathways were upregulated in different cell types (Table S10). In endothelial cells, positive regulation of cell migration (GO:0030335) genes, in Leydig cells, cholesterol biosynthesis (GO:0006695), in macrophages, peptide antigen assembly with MHC protein complex (GO:0002501), and in myoid cells, regulation of apoptotic process (GO:0042981) were the top enriched genes that were upregulated. These results indicated that despite downregulation of RP genes, different pathway genes were differentially upregulated in NOA somatic cells compared to normal somatic cells of the testis.

We also compared RP gene expression in the somatic and germ cells of a patient with retrograde oligozoospermia with secondary infertility because of ejaculatory dysfunction. While the RP gene expression patterns in the somatic cells were largely similar to NOA (Figure S4), in some of the available germ cells, namely, spermatids and sperm cells, RP gene expression was not significantly different from that of the normal cells. In the spermatogonial stem cells, 9 RP gene expression values were downregulated compared to normal cells (Figure S4, Table S9). In addition, in Klinefelter syndrome, RP gene expression in somatic cells was similar to that of somatic cells of a normal testis, with only *RPS26* being upregulated in all somatic cells (Figure S5, Table S9).

### Disrupted RP gene expression characterises Diamond-Blackfan Anaemia haematopoiesis

We wanted to compare RP gene expression changes during spermatogenesis, a meiotic process, with erythropoiesis, a mitotic process, which is the differentiation of haematopoietic stem cells to mature erythrocytes. The RP gene expression data of many erythroid progenitor cells in the bone marrow tissue of a normal patient and two patients with DBA were arranged in a pseudotime line, with the progenitor cells with high expression of *KIT* as the starting point (Figure S6)^55^.

The clusters of RP gene expression patterns during erythropoiesis, identified using DPGP analysis, found that during normal erythropoiesis, 72 out of 76 RP genes cluster together and follow the same pattern with lower expression as the erythrocytes mature (Figure 5 A-B). Only 4 RP genes, namely, *RPL17, RPL26, RPL15,* and *RPS17*, formed a separate cluster with a different expression pattern (Figure 5A-B). When the donors were analysed separately, we predominantly found these two patterns, and 64 out of 76 RP genes commonly showed a pattern similar to the first cluster when the donors were combined (Figure S7).

**Figure 5.**
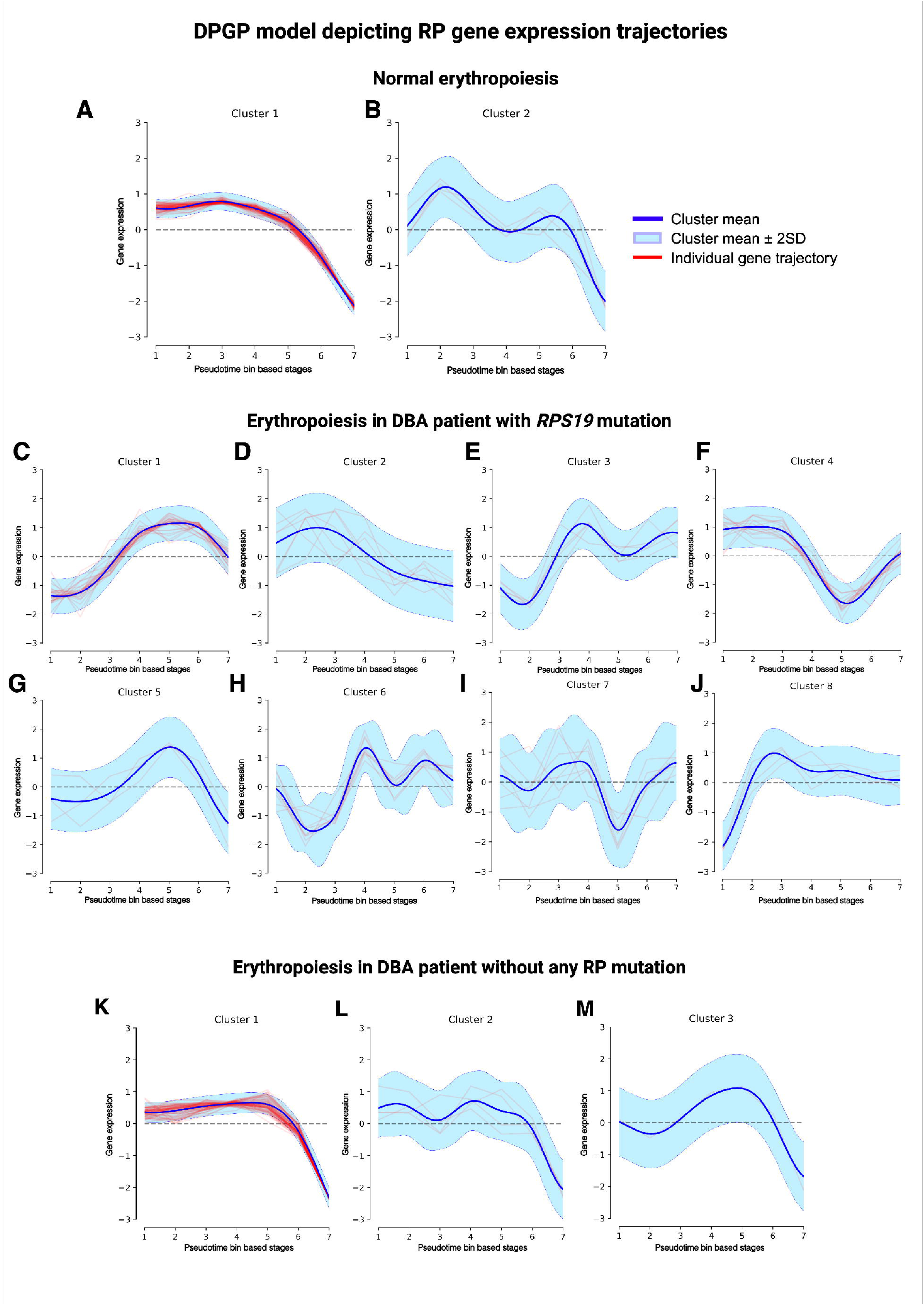
**Temporal clustering reveals distinct waves of RP gene expression during erythropoiesis**. The DPGP model-based trajectories of RP gene expression profiles along erythropoiesis (x-axis). The y-axis represents the log-normalised ScTransform corrected gene expression values of RP genes, and the x-axis represents different pseudotime bins along the erythropoiesis trajectory. All genes were grouped based on their shared temporal expression dynamics across the pseudotime trajectory established in panel (A). Overview of 14 distinct clusters: (A-B) normal erythropoiesis, (C-J) erythropoiesis in the DBA patient with *RPS19* mutation, (K-M) the patient with clinical DBA without RP mutation. Each small plot displays the normalised expression (y-axis) of individual RP genes within that cluster (red lines) as a function of pseudotime (x-axis). The “light blue zone” along each line in the plots represents the standard deviation of gene expression for the individual RP genes within that cluster, around the mean expression profile (blue line). Created in https://BioRender.com.

When we analysed RP gene expression patterns in a DBA patient with *RPS19* L18P mutation, we found complete dysregulation of RP gene expression patterns with eight clusters (Figure 5C-J). Interestingly, the mutated *RPS19* followed a completely opposite expression pattern from that of normal individuals, where the expression of this RP gene increased along the erythropoiesis progression. In the second DBA patient with no identified RP gene mutation but with clinical symptoms, 71 out of 76 RP genes followed the expression pattern observed in the case of normal, and the remaining 5 RP genes belonged to a different cluster with a similar expression pattern (Figure 5K-M). The RP genes belonging to different clusters in normal and DBA erythrocytes are given in Table S11. Further, in order to understand the change of expression patterns of RP gene upstream regulators, we looked at how *GATA1* and *SPI1* expression change along the pseudotime in both normal and DBA patients (Figure 6). *GATA1* expression increased during erythropoiesis progression, similar to *RPS19* in DBA with *RPS19* mutation (Figure 6B). In contrast, it decreased along pseudotime in normal erythropoiesis (Figure 6A) and in DBA patients without RP mutation (Figure 6C), while *SPI1* expression remained unchanged in all three cases. These results indicated that the expression patterns of the RP genes during erythropoiesis were similar to the changes observed in *GATA1*. In case of DBA with RP mutations, a dysregulation was observed in both RP genes and *GATA1*.

**Figure 6.**
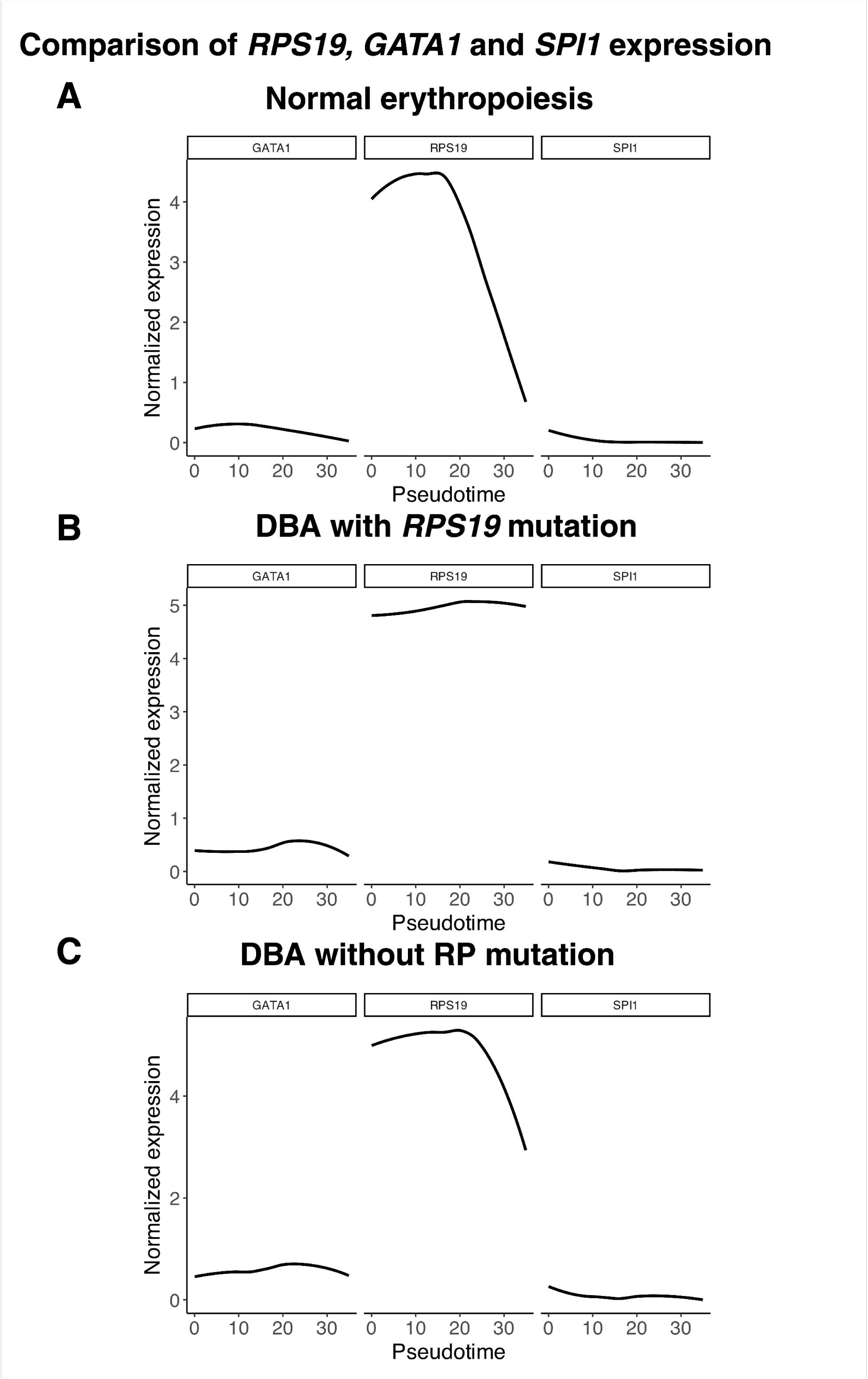
**Differential Expression of *RPS19*, *GATA1*, and *SPI1* During Erythroid Differentiation in Normal and DBA**. LOESS smoothed curve representing normalised expression levels of *GATA1*, *RPS19*, and *SPI1* plotted across pseudotime, representing the erythroid differentiation process in (A) normal erythropoiesis, (B) DBA with an *RPS19* mutation, and (C) DBA without any RP mutation. The log-normalised ScTransform corrected gene expression values (y-axis) are plotted along the pseudotime along the erythropoiesis trajectory (x-axis). The gene expression in each cell along the pseudotime is plotted as a LOESS smoothed curve. Created in https://BioRender.com.

## DISCUSSION

Specialised ribosomes, identified in bacteria to humans^76,33,18,5^, have the ability to preferentially translate specific transcripts to elicit environmental or tissue-specific functions^20,18,21,17^. Distinct functions of specialised ribosomes arises from many mechanisms^4,77,78^, including differential stoichiometry of RP genes, which can be indicated by the heterogeneous expression of these genes. In humans, the tissue-specific RP gene expression heterogeneity has been shown in bulk RNA sequencing datasets^23,31,29^. In this study, by analysing 161 cell types from 15 healthy tissues, we establish that each RP gene expression level is highly variable depending on the tissue and cell type within the tissue.

Despite being in the same tissue, somatic and germ cells in the adult testis had divergent RP gene expression patterns. The germ cells, namely, spermatogonial stem cells, differentiating spermatogonia, early and late primary spermatocytes, round and elongated spermatids, and sperm, showed high RP gene expression variance, unlike expression in somatic cells. The high variance in RP gene expression, which is specific to each stage, could indicate the presence of specialised ribosome pools that are active during various stages of spermatogenesis. This stage-specific specialisation of ribosomes may suggest distinct functions at different stages of spermatogenesis. These results align with a previous study where the role of ribosomes is stage-specific, such that in spermatocytes, ribosomes have active translational functions, whereas in round spermatids, they are largely stalled and associated with nonsense-mediated decay^79^.

Previous studies have shown that RP genes are crucial for sperm development in multiple species^10,22,73,80–83^. Interestingly, in *Drosophila*, *RPS25* role was found to be stage-specific, vital for the elongation stage but not the early stages of spermatogenesis^84^. Similarly, we observed that *RPS25* expression decreases during the early stages of spermatogenesis and increases during the stage when spermatids begin to elongate. Our analysis revealed that *RPL3* and *RPS6*, the two ribosomal protein genes previously identified as playing a role in sperm motility, were expressed in sperm^73^. Thus, differential RP gene expression, which could lead to differential stoichiometry and therefore, heterogeneous specialised ribosomes, has been shown to preferentially translate specific mRNA, resulting in specific functions^20,34,25^.

Altered RP gene expressions in male infertility have been reported at a bulk level in asthenozoospermia^85^ and NOA^86,87^. In NOA, it was shown that *RPS3* and *RPS2* have decreased expression^87^. Our analysis of NOA confirmed these results and distinguished that this decreased expression is cell-type specific, *RPS2* was downregulated in somatic cells, and *RPS3* was downregulated only in the endothelial cells. Further, we observed many RP genes downregulated in the somatic cells of individuals with male infertility. Given the crucial supportive role of these somatic cells in spermatogenesis, their altered RP landscape could impair their functionality, contributing to the infertility phenotype^74,75^. This cell-type-specific insight into RP dysregulation offers more granular targets for understanding spermatogenesis and male fertility than bulk tissue analyses.

While in spermatogenesis, RP gene expression shows variable stage-specific patterns, in erythropoiesis, no such patterns were identified, indicating that RP genes had similar expression in each stage. Further, we observed that in erythropoiesis, RP gene expression reduces as the differentiation progresses. Similarly, previous studies have shown that protein levels of RP genes decrease with erythroid differentiation^88^, and ribosome biogenesis is reduced as erythroid differentiation progresses^89–91^. Additionally, in DBA patients with *RPS19* mutation, RP gene expression patterns are dysregulated compared to normal erythropoiesis. Previous studies have shown that *RPS19* plays a critical role in regulating ribosome biogenesis, and its deletion leads to defective ribosome formation^92^ and the effects of *RPS19* mutations were very specific to erythroid differentiation^93^.

The mRNA and protein levels of RP genes show a higher correlation with each other compared to other gene sets^31,94^. This suggests that the changes in the transcript levels of RP genes observed in this study may also correspond to changes at the protein level across different cells, tissues, and pathologies. This implies that there may be varying protein-level stoichiometric changes in different cell types and tissues. These findings highlight the diverse functions of RP genes and specialised ribosomes, which contribute to the specificity of cellular functions and help identify novel signatures of disease pathology.

## Supporting information

Supplementary Results

## ACKNOWLEDGEMENTS

We thank Devansh Sanghvi, Srijith Sasikumar, Veerendra Gadekar, and Venkatesh Kamaraj (IIT Madras) for their assistance with computational analysis. We thank Kedar Natarajan and Manikandan Narayanan for critically reviewing the manuscript. We appreciate the members of the Systems Genetics Lab, the Centre for Integrative Biology and Systems Medicine (IBSE) and Wadhwani School of Data Science and AI (WSAI) for their inputs and insightful discussions.

## FUNDING

A.M. was supported by a fellowship from the Council of Scientific & Industrial Research (CSIR) Fellowship (09/084(0774)/2020-EMR-I), an International Immersion Experience grant from Office of Global Engagement, IIT Madras, a Women Leading Science grant from IIT Madras (SB/25-26/0034/BT/IITM/008752). A.M. was also supported by a grant from the Centre for Integrative Biology and Systems Medicine (IBSE), IIT Madras (BIO/18-19/304/ALUM/KARH) and Excelra Knowledge Solutions Private Limited (CR/22-23/0026/BT/EXCE/008752) to H.S. A.D.J. was supported by the Singapore Ministry of Health’s National Medical Research Council Clinician Scientist Award (MOH-000715-00). Work in the A.D.J. laboratory is funded by the core grant from the Cancer Science Institute of Singapore, National University of Singapore, through the National Research Foundation Singapore and the Singapore Ministry of Education under its Research Centres of Excellence initiative.

## AUTHOR CONTRIBUTIONS

H.S. and A.M. conceptualised the project; A.M. developed the computational pipeline; AM analysed and interpreted the data; C.T. and A.D.J. contributed to the analysis; A.M. and H.S. wrote the manuscript; all authors commented and edited the manuscript.

## DATA AND CODE AVAILABILITY

The codes used in this paper are available on GitHub (https://github.com/HimanshuLab/HCA-scRNAseq-RP-genes).

## DECLARATION OF INTERESTS

The authors declare no competing interests.

## SUPPLEMENTARY INFORMATION

Supplementary information, including results, figures, and tables, accompanies this manuscript.

## Notes

### Competing Interest Statement

The authors have declared no competing interest.

