## Supplementary Results for "A single-cell atlas of ribosomal protein heterogeneity across human tissues reveals phenotypes of biological and clinical significance"

### ^4^ Cancer Science Institute of Singapore, National University of Singapore, Singapore, Singapore

^5^ Department of Haematology-Oncology, National University Health System, Singapore, Singapore

^6^ NUS Centre for Cancer Research, Yong Loo Lin School of Medicine, National University of Singapore, Singapore, Singapore

^7^ Department of Medicine, Yong Loo Lin School of Medicine, National University of Singapore, Singapore, Singapore

#### **SUPPLEMENTARY RESULTS**

#### **Comparison of different donors in the testis for the heterogeneity of RP gene expression**

#### To investigate the variability in RP gene expression among different donors in the testis, we calculated the IOD values for each donor by combining two replicates. Our observations revealed that most RP genes exhibited consistently high IOD values across all three donors. The hierarchical clustered heatmap illustrates the mean expression across cell types and the IOD values of RP genes for all three testis donors (Figure S1A, B). Notably, donor 2 demonstrated higher RP gene expression levels than the other two donors. When comparing IOD values, 30 RP genes exhibited high IOD across all three donors, while some RP genes displayed slight variations in their IOD values. This could be due to the differences in the number of constituent cell types in each donor. The RP genes with high IOD (above the 90th percentile) for each donor in the testis are displayed in Figure S1C. The differences in IOD between donors can be attributed to variations in the composition of testis cell types for each case (Figure S1D).

#### **Ribosomal protein genes are distinct from other random and ubiquitous genes in terms of their gene expression variance within the tissue.**

A list of ubiquitously expressed genes was obtained from Gu et al.^1^ From this list, pathways with approximately 100 genes were selected for analysis, including cellular respiration, chromatin remodelling, DNA repair, endosomal transport, phosphorylation, RNA metabolism, and RNA polymerase transcription. For each chosen pathway, the Seurat object of each tissue was subset to include only genes from that pathway. The mean and variance of each gene were calculated throughout the tissue to calculate the IOD. Levene’s test was performed to determine whether the distribution of RP gene expression IOD differed significantly from other genes, followed by a pairwise comparison to identify the significantly different pairs. This comparison showed that the RP genes had a higher range of IOD than the ubiquitously expressed gene sets for almost all tissues. The Levene’s test to test the significant difference between the data distribution between these gene sets showed that RP genes had a significantly different distribution compared to all other ubiquitous gene sets studied in the testis. While in many tissues like the bile duct, oesophagus, liver, marrow, spleen and stomach, the range of IOD shown by RP genes was similar to the cellular respiration genes but different from other gene sets. The pairwise Levene’s test results for each tissue are shown in Table S12. This result shows that not all RP genes are similarly variable, and the range of variability within a tissue is much higher than it is with other genes.

To establish a baseline for comparison, a similar analysis was conducted on 20 sets of 76 randomly selected genes, excluding RP genes, which are different for each tissue. The list of random genes used for each tissue is given in Table S13. The IOD of these random gene sets was calculated and analysed using the same approach as for the pathway-specific gene sets. To account for the higher mean of RP genes compared to other gene sets and to enable comparisons between them, we calculated the IOD. In the testis, bladder, and muscles, the RP genes showed increased distribution of the IOD, which was found to be significantly different from random gene sets using Levene’s test (Table S14). However, in other tissues, analogous to the previous result of IOD values of RP genes, they were not very high and significantly different from other random genes, except for a few. These results demonstrate that RP genes exhibit varying degrees of expression variability across different tissues, with unique expression patterns particularly distinctive to RP genes in certain tissues. Notably, the testis shows a significantly higher variability in RP gene expression, a pattern not observed in other gene sets. This distinct variability highlights the importance of studying RP gene expression specifically within the testis.

**SUPPLEMENTARY FIGURES**

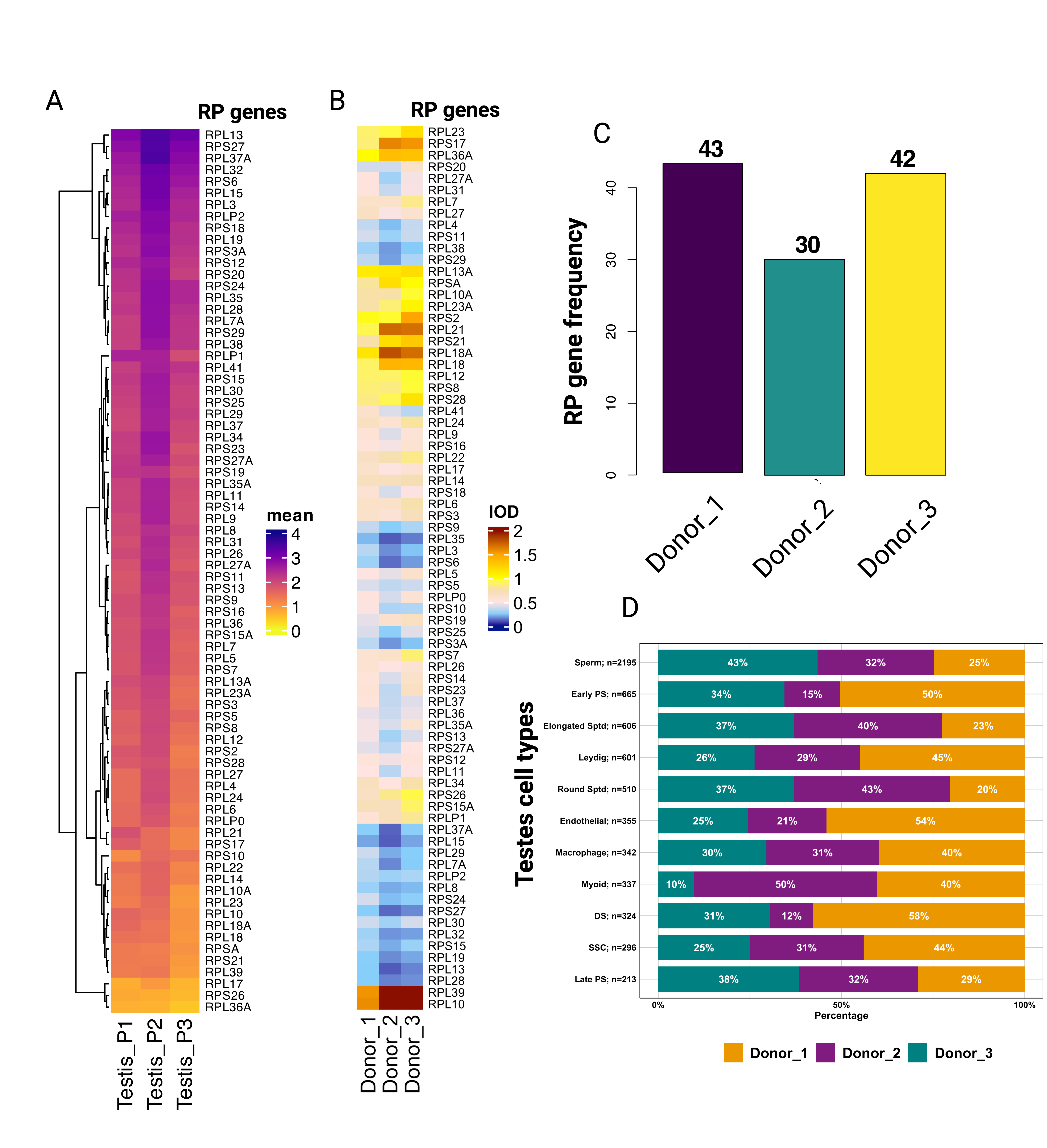

**Figure S1. RP heterogeneity in different donors of the testis**

(A) Heatmap depicting the mean normalised expression of 76 cytoplasmic RP genes (rows) across 3 different donors of testis (columns). There are three donors: donor 1, donor 2, donor 3 (x-axis) and 76 cytoplasmic RP genes (y-axis). Colour intensity scale bar (0 to 4) corresponds to the average SCTransform-normalised expression level of RP genes within each tissue. RP genes are hierarchically clustered. (B) Heatmap depicting the IOD values of gene expression of 76 cytoplasmic RP genes (rows) across 3 different donors of testis (columns). There are three donors, namely, donor 1, donor 2, and donor 3. The colour scale bar represents the IOD (scale 0 to 2), calculated as variance divided by the mean of normalised expression for RP genes within each tissue. The x-axis lists the donors, and the y-axis the cytoplasmic RP genes. (C) Histogram showing the number of RP genes that exhibit a high IOD value (above the 90^th^ percentile) in each donor of the testis. The y-axis represents the count of RP genes having high IOD, and the x-axis lists different donors of tests, namely, Donor_1, Donor_2, and Donor_3. (D) Bar plot showing the distribution of different donors in each cell type. The y-axis gives the cell type of the testis, and the x-axis depicts the percentage of each cell type in each donor. Each donor is depicted in different colours, Donor_1 (orange), Donor_2 (violet), and Donor_3 (green), and the percentage of donors in each cell type is mentioned in the graph. The total number of cells in each cell type and the respective cell types are shown on the y-axis. Created in <https://BioRender.com>

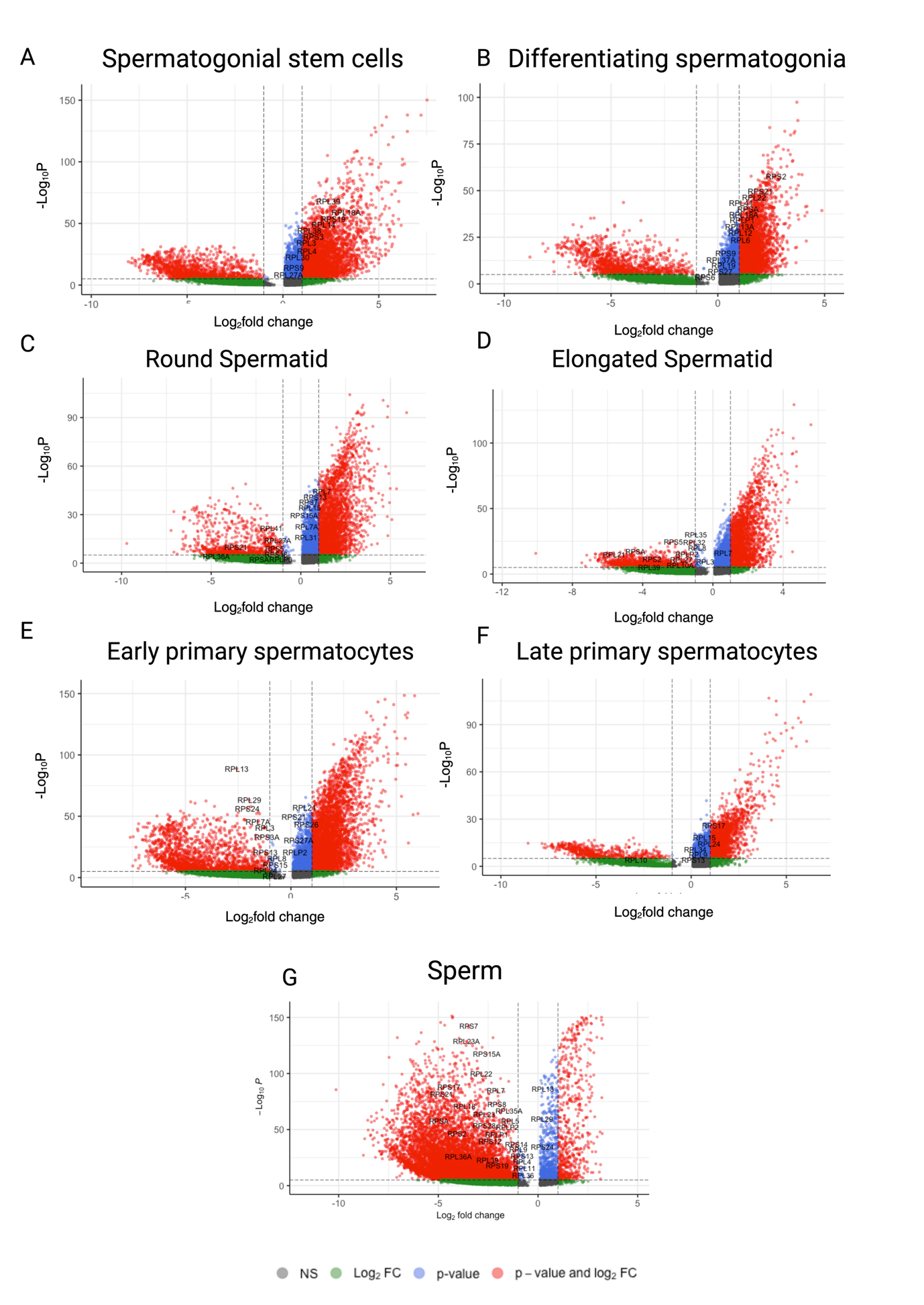

**Figure S2.** **Differential Expression of RP Genes During Spermatogenesis**.

Volcano plots illustrating differential gene expression changes in various spermatogenic cell types compared with one another. The x-axis of each plot represents the log_2_fold change, while the y-axis indicates the −log_10_(p-value). Each point in this plot depicts a gene. Genes are coloured based on statistical significance: NS (not significant, grey), significant by log_2_fold change only (green), significant by p-value only (blue), and significant by both p-value and log_2_fold change (red). The individual panels represent the differential expressed genes in individual cell types compared with other germ cell types: (A) Spermatogonial stem cells, (B) Differentiating spermatogonia, (C) Early primary spermatocytes, (D) Round Spermatid, (E) Late primary spermatocytes, (F) Elongated Spermatid, (G) Sperm. Created in <https://BioRender.com>

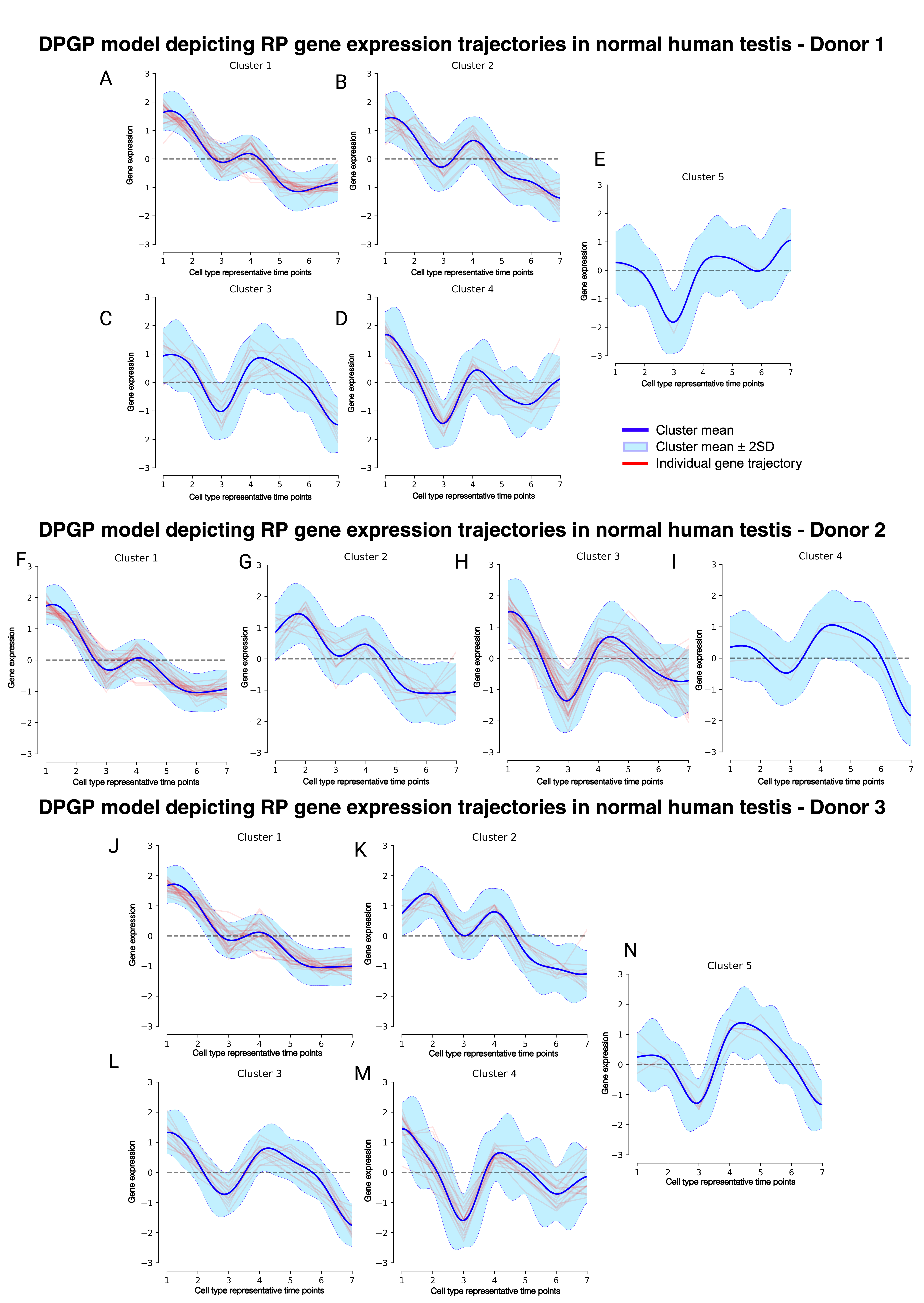

**Figure S3. The DPGP model depicts RP gene expression trajectories in normal human testis from three donors.**

The DPGP model-based trajectories of RP gene expression profiles along spermatogenesis (x-axis) of 14 distinct gene clusters (Clusters 1-5). The y-axis represents the log-normalised ScTransform corrected gene expression values of RP genes. The x-axis represents the cell-type states of spermatogenesis at different representative time points: Spermatogonial stem cells (1), differentiating spermatogonia (2), early primary spermatocytes (3), late primary spermatocytes (4), round spermatids (5), elongated spermatids (6), sperms (7). Each small plot displays the normalised expression (y-axis) of individual RP genes within that cluster (red lines) as a function of pseudotime (x-axis). The “light blue zone” along each line in the plots represents the standard deviation of gene expression for the individual RP genes within that cluster, around the mean expression profile (blue line). (A-E) Gene expression trajectories for Donor 1, showing different clusters: (A) Cluster 1, (B) Cluster 2, (C) Cluster 3, (D) Cluster 4, and (E) Cluster 5. (F-I) Gene expression trajectories for Donor 2, showing different clusters: (F) Cluster 1, (G) Cluster 2, (H) Cluster 3, and (I) Cluster 4. (J-N) Gene expression trajectories for Donor 3, showing different clusters: (J) Cluster 1, (K) Cluster 2, (L) Cluster 3, (M) Cluster 4, and (N) Cluster 5. Created in <https://BioRender.com>

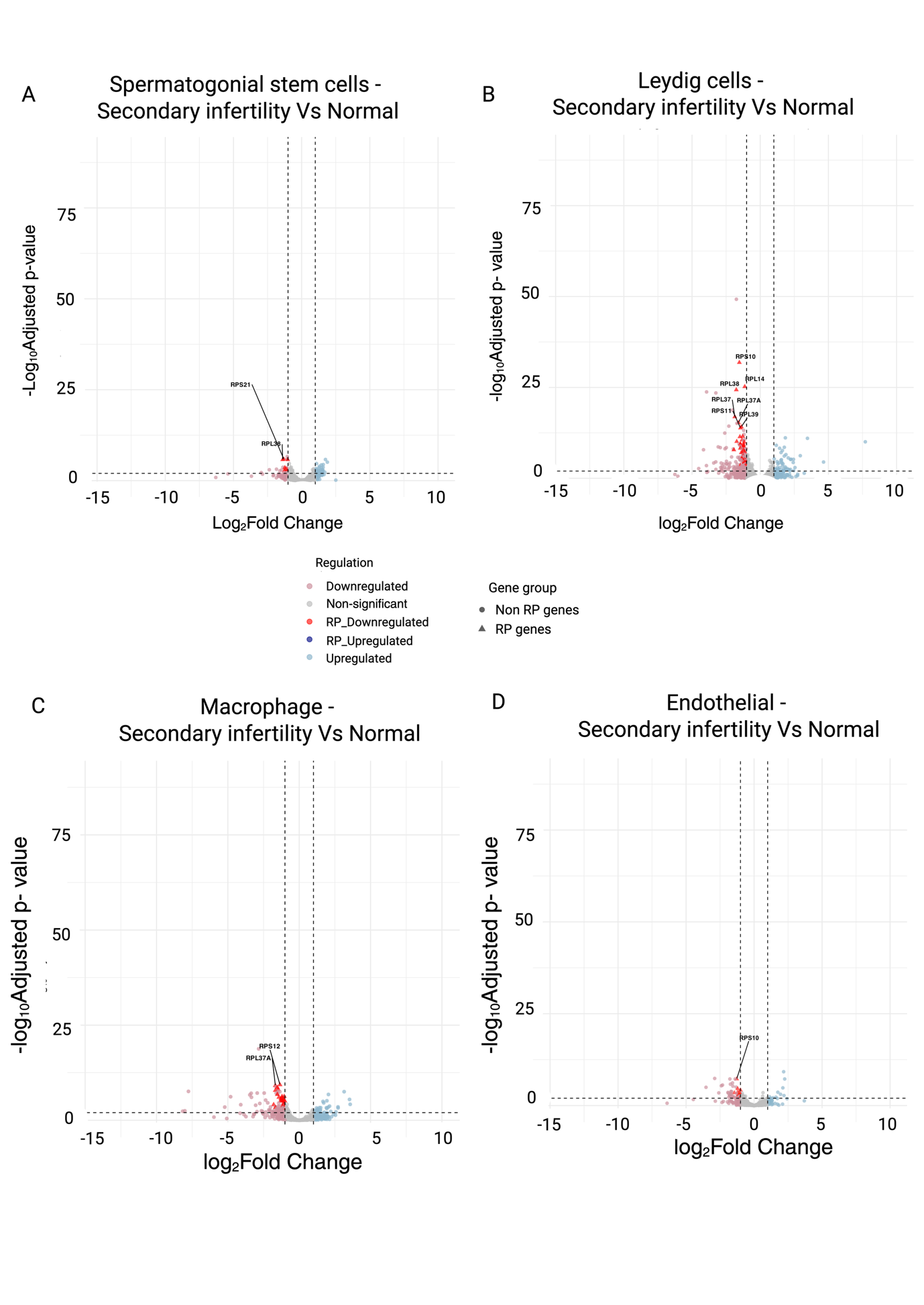

**Figure S4. Differential RP gene expression in testicular cell types of individuals with retrograde oligospermia compared to normal controls.**

Volcano plots demonstrating differential gene expression in the pseudobulk aggregation in testicular cell types when comparing samples from individuals with retrograde oligospermia (secondary infertility) to normal controls: (A) Spermatogonial stem cells, (B) Leydig cells, (C) Macrophages, (D) Endothelial cells. In these plots, each point signifies a gene. The x-axis displays the log_2_ fold change, while the y-axis represents the -log_10_ (p-value). Genes showing statistically significant upregulation or downregulation are highlighted, with RP genes that are significantly upregulated “RP_Upregulated” (dark blue), RP genes that are significantly downregulated “RP_Downregulated” (dark red), Other, non-RP genes that are significantly upregulated “Upregulated” (light blue), Other, non-RP genes that are significantly downregulated “Downregulated” (light red), Non-significant genes “Non-significant” (grey). RP genes are indicated as triangles, and all other genes are indicated as circles. Created in <https://BioRender.com>

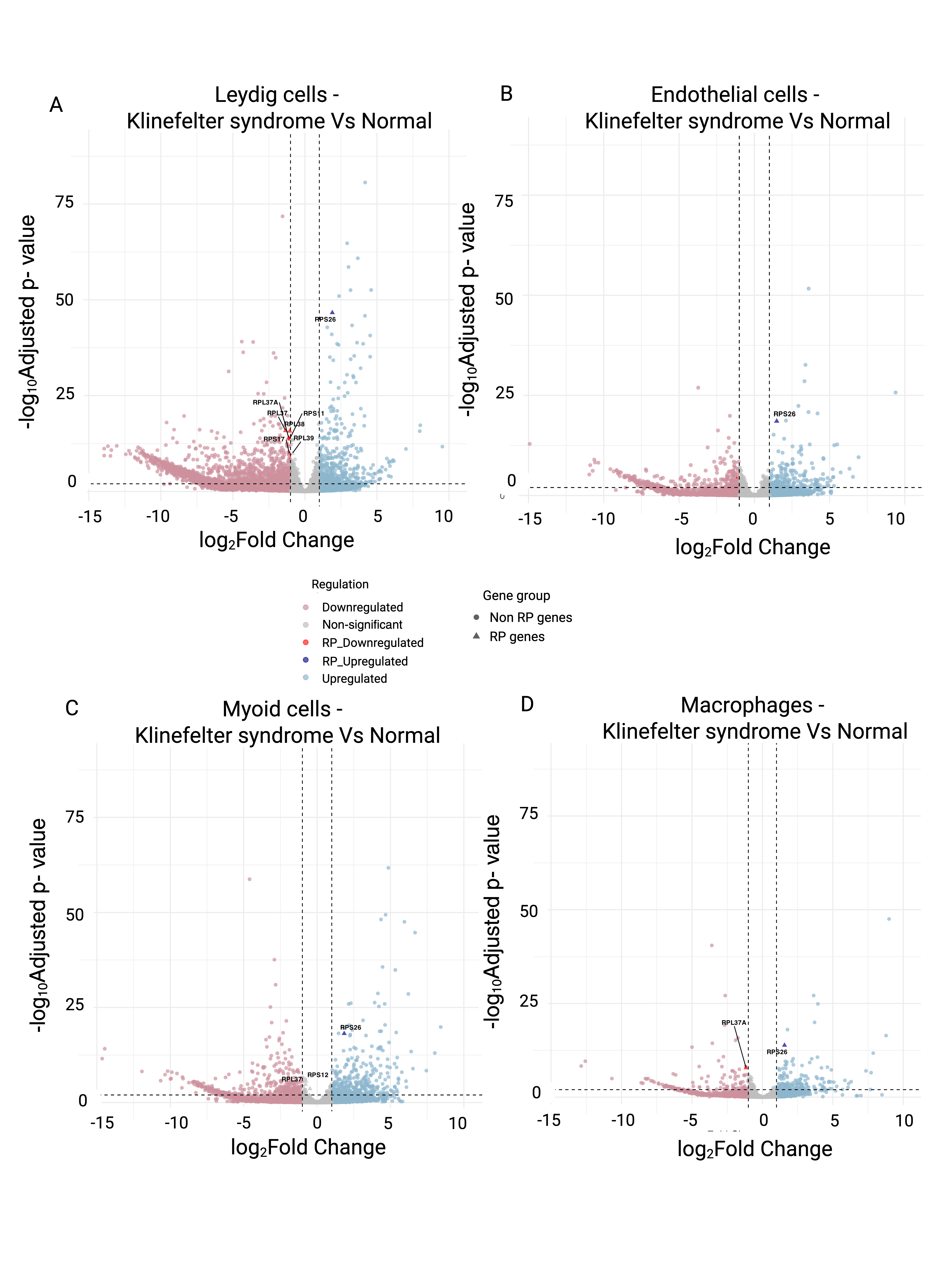
**Figure S5. Differential RP gene expression in testicular cell types of individuals with Klinefelter syndrome compared to normal controls.**

Volcano plots demonstrating differential gene expression in the pseudobulk aggregation in testicular cell types when comparing samples from individuals with Klinefelter syndrome to normal controls: (A) Leydig cells, (B) Endothelial cells, (C)Myoid cells, (D) Macrophages. In these plots, each point signifies a gene. The x-axis displays the log_2_ fold change, while the y-axis represents the -log_10_ (p-value). Genes showing statistically significant upregulation or downregulation are highlighted, with RP genes that are significantly upregulated “RP_Upregulated” (dark blue), RP genes that are significantly downregulated “RP_Downregulated” (dark red), Other, non-RP genes that are significantly upregulated “Upregulated” (light blue), Other, non-RP genes that are significantly downregulated “Downregulated” (light red), Non-significant genes “Non-significant” (grey). RP genes are indicated as triangles, and all other genes are indicated as circles. Created in <https://BioRender.com>

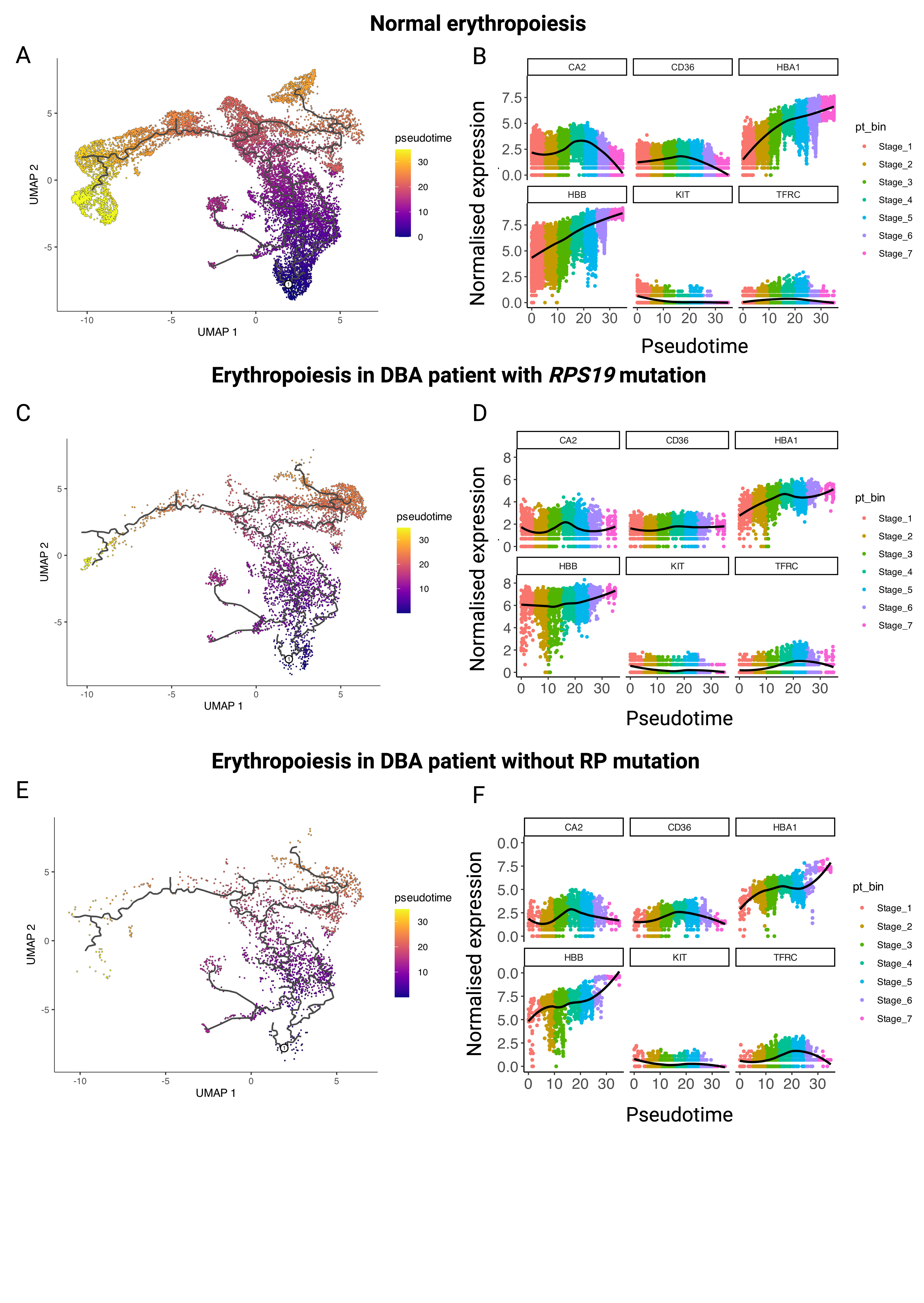

**Figure S6. Pseudotime Trajectory of erythropoiesis and associated Gene Expression Dynamics.**

UMAP plots and marker gene expression trajectories for normal erythropoiesis (A, B), DBA with *RPS19* mutation (C, D) and DBA without RP mutation (E, F). Panels A, C, and E show UMAP plots of erythropoiesis progression, with cells coloured by 'pseudotime', indicated by the colour bar ranging from purple (low pseudotime) to yellow (high pseudotime). The black lines overlaid on the UMAPs represent the inferred pseudotime trajectory or principal curves. Panel B, D, and F show scatter plots of scTransform normalised expression (y-axis) versus pseudotime for several genes: *CA2, CD36, HBA1, HBB, KIT,* and *TFRC*. The x-axis represents pseudotime, aligning with the pseudotime calculated in the UMAPs (Panels A, C, E). Each point in the scatter plot represents a cell. The cells are coloured by pseudotime bin, indicating different stages (Stage_1 to Stage_7), which are discreet groupings of cells along the pseudotime axis. A black line (LOESS smoothing curve) is overlaid on each gene's expression profile, showing the trend of gene expression as pseudotime progresses. Created in <https://BioRender.com>

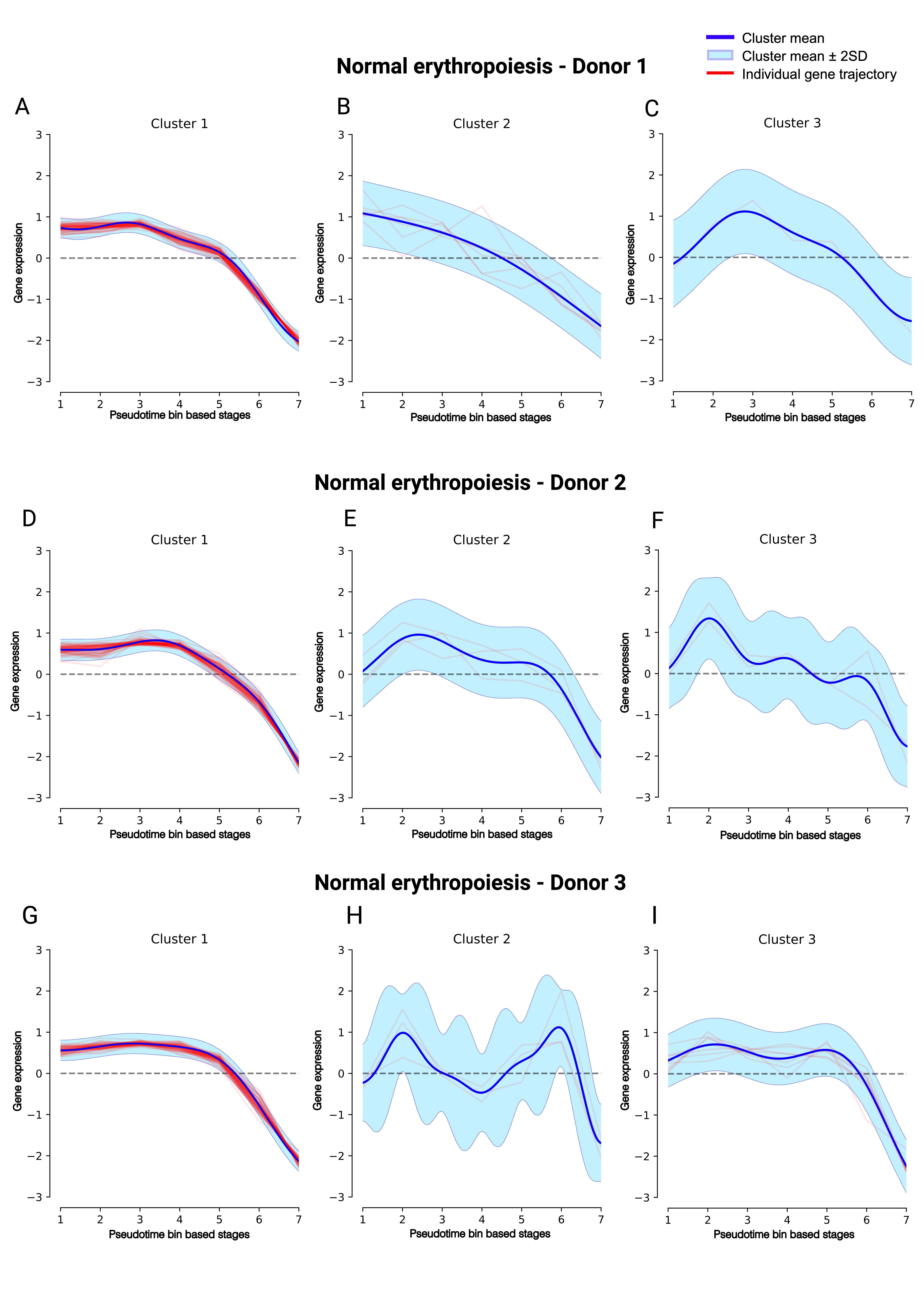
**Figure S7.** **RP gene expression changes in erythropoiesis have consistent patterns in different donors.**

The DPGP model-based trajectories of RP gene expression profiles along erythropoiesis (x-axis). The y-axis represents the log-normalised ScTransform corrected gene expression values of RP genes, and the x-axis represents different pseudotime bins along the erythropoiesis trajectory. All genes were grouped based on their shared temporal expression dynamics across the pseudotime trajectory established in panel (A). Overview of 8 distinct clusters: (A-C) donor 1, (D-F) donor 2, (G-H) donor 3. Each small plot displays the normalised expression (y-axis) of individual RP genes within that cluster (red lines) as a function of pseudotime (x-axis). The “light blue zone” along each line in the plots represents the standard deviation of gene expression for the individual RP genes within that cluster, around the mean expression profile (blue line). Created in <https://BioRender.com>

**SUPPLEMENTARY TABLES**

**Table S1:** Overview of the datasets used in this study for the 15 tissues and the preprocessing parameters.

**Table S2:** Mean expression values for RP genes across the constituent cell types of 15 tissues and the three donors of the testis.

**Table S3:** IOD values for RP genes across the constituent cell types of 15 tissues and the three donors of the testis.

**Table S4:** Cell-specific marker genes for cell identity assignment.

**Table S5:** Differentially expressed genes identified across cell types in various tissues and across various tissues in common cell types.

**Table S6:** Commonly differentially expressed genes identified across the cell types of adult testis across different donors.

**Table S7:** Commonly differentially expressed genes identified across the cell types of infant testis across different donors.

**Table S8:** Clustered RP genes in testis identified by DPGP analysis of individual and combined donor samples.

**Table S9:** Differentially expressed genes in various testis cell types of non-obstructive azoospermia, retrograde oligospermia, and Klinefelter syndrome compared to healthy testicular cells.

**Table S10:** Gene ontology analysis of differentially expressed genes in NOA testes.

**Table S11:** Clustered RP genes in normal erythropoiesis identified by DPGP analysis of individual and combined donor samples.

**Table S12:** Pairwise comparison of RP genes with ubiquitous genes using Levene’s test in different tissues.

**Table S13:** List of random gene sets for each tissue compared with RP gene IOD.

**Table S14:** Pairwise comparison of RP genes with random gene sets using Levene’s test in different tissues.
